# A systematic search for RNA structural switches across the human transcriptome

**DOI:** 10.1101/2023.03.11.532161

**Authors:** Matvei Khoroshkin, Daniel Asarnow, Albertas Navickas, Aidan Winters, Johnny Yu, Simon K. Zhou, Shaopu Zhou, Christina Palka, Lisa Fish, K. Mark Ansel, Yifan Cheng, Luke A. Gilbert, Hani Goodarzi

## Abstract

RNA structural switches are key regulators of gene expression in bacteria, yet their characterization in Metazoa remains limited. Here we present SwitchSeeker, a comprehensive computational and experimental approach for systematic identification of functional RNA structural switches. We applied SwitchSeeker to the human transcriptome and identified 245 putative RNA switches. To validate our approach, we characterized a previously unknown RNA switch in the 3’UTR of the RORC transcript. *In vivo* DMS-MaPseq, coupled with cryogenic electron microscopy, confirmed its existence as two alternative structural conformations. Furthermore, we used genome-scale CRISPR screens to identify *trans* factors that regulate gene expression through this RNA structural switch. We found that nonsense-mediated mRNA decay acts on this element in a conformation-specific manner. SwitchSeeker provides an unbiased, experimentally-driven method for discovering RNA structural switches that shape the eukaryotic gene expression landscape.

## INTRODUCTION

Gene expression is regulated at the RNA level in all kingdoms of life. The two oldest groups of RNA-based regulatory mechanisms are ribozymes (catalytically active RNA molecules) and RNA structural switches (or riboswitches). RNA switches are regulatory elements that control gene expression by direct binding of a small-molecule ligand or other *trans*-acting factor. In bacteria, RNA switches are one of the most widely observed mechanisms for gene expression control. Classic bacterial riboswitches bind small molecule ligands directly, inducing a conformational change, leading to modulation of expression of the host transcript (Serganov and Nudler 2013). The search for such ligand-binding riboswitches in eukaryotes has had limited success to date. Just two human examples are known: the RNA switch in VEGFA and the m6A modification-based switches (Liu et al. 2015; Ray et al. 2009), and it remains unclear how widely this regulatory mechanism impacts gene expression in higher eukaryotes despite its ubiquity in other domains of life. Here, we introduce SwitchSeeker, a systematic, computational and experimental framework for unbiased discovery of RNA structural switches across any transcriptome.

While several RNA switch detection software packages have been developed, most identify new switch sequences based on their homology to one of the 40 known RNA switch families (Kalvari et al. 2021). Several tools for *de novo* discovery of RNA switches have been developed; however, their predictions often lack experimental verification of both structure and function (Barsacchi et al. 2016; Manzourolajdad and Arnold 2015). Therefore, there is a need for scalable methods of detecting eukaryotic RNA switches and assessing the extent to which they carry out regulatory functions in gene expression control. Our approach relies on integrating multiple computational and experimental methods, where RNA switches are first predicted *in silico*, then structurally and functionally characterized *in vivo*, which in turn informs the next iteration of *in silico* predictions. First, we developed a computational model for *de novo* RNA switch detection, named SwitchFinder – a component of our SwitchSeeker framework. We showed that SwitchFinder identifies RNA switches from novel families with higher accuracy than the existing models. We applied SwitchFinder to the human transcriptome to select putative RNA switches. We then used massively parallel assays *in vivo* to interrogate both the structure and function of these candidate elements. By iteratively improving the SwitchFinder predictions with experimental data, we reported ∼250 high-confidence and functional RNA structural switches.

Finally, we selected the top scoring switch, located in the 3’UTR of the RORC transcript, for further analysis and dissection. We used DMS-MaPseq structural probing and single-particle cryogenic electron microscopy (cryo-EM) to confirm that the predicted switch populates alternate molecular conformations. We then performed genome-scale CRISPR-interference (CRISPRi) screens, which revealed that one of the two conformations reduces gene expression through activation of the non-canonical nonsense-mediated decay (NMD) pathway. Taken together, our framework provides new insights into the role of RNA structural switches in shaping the transcriptome in human cells, and outlines a broader approach for future characterization of RNA switches regulating eukaryotic gene expression control across cell types.

## RESULTS

### Systematic annotation of human RNA structural switches

We first set out to identify RNA sequences that exist in multiple structural conformations. For this, we developed SwitchFinder, a tool that predicts whether an RNA sequence contains putative RNA structural switches. We designed SwitchFinder to satisfy several criteria: first, it should predict if a given RNA sequence contains a potential RNA switch and suggest the two mutually exclusive folding conformations (examples are shown in Fig. 1A, Suppl. Fig. 1A). Secondly, it should be able to effectively capture a more generalizable definition of RNA switches in order to find instances beyond the 40 known RNA switch families (Kalvari et al. 2021) (for example, the human RNA switch in VEGFA gene, Fig. 1A <u>(Ray et al. 2009)</u>). Thirdly, it should allow for seamless integration of experimental data to improve predictions. This is especially important as mRNA secondary structure in the cell is shown to be highly dynamic (Mortimer, Kidwell, and Doudna 2014; Rouskin et al. 2014) and compartment-dependent (Sun et al. 2019); therefore, the predictions can be greatly improved with *in vivo* secondary structure probing data.

**Fig 1:**
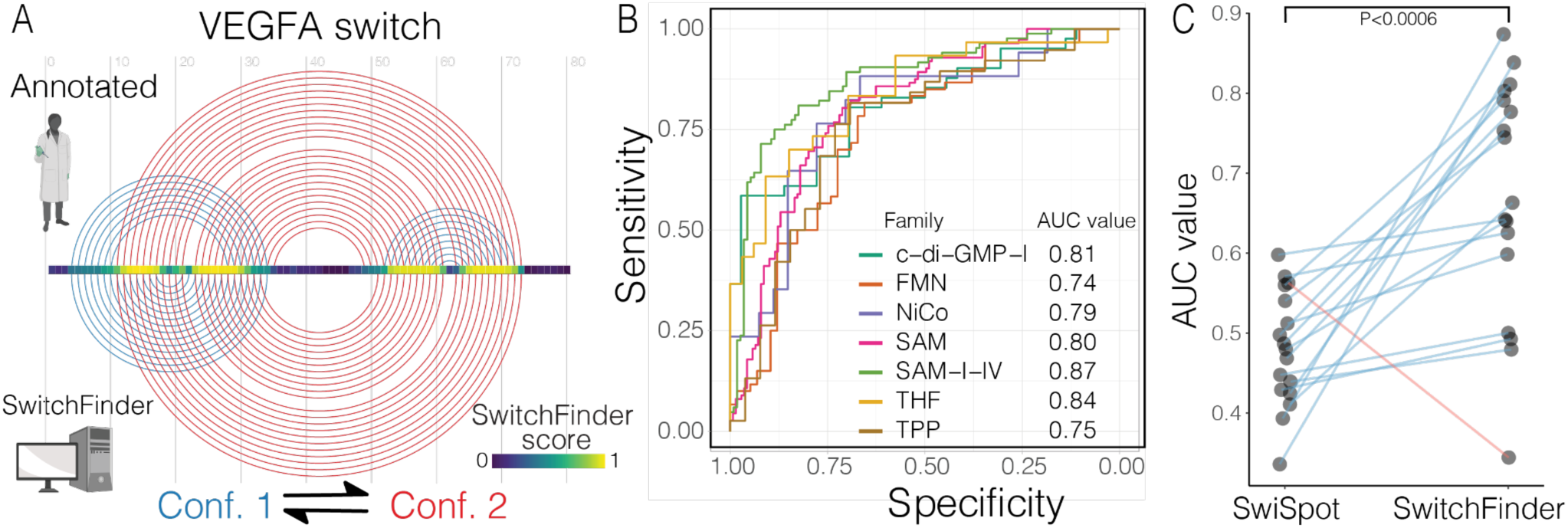
SwitchFinder identifies candidate RNA switches in the human genome. (A) Example of SwitchFinder locating the RNA switch within the VEGFA mRNA sequence. Top: arc representation of the RNA base pairs that change between the two conformations of VEGFA RNA switch, as in (Ray et al. 2009). The two conformations are shown in red and blue, respectively. Bottom: the two conformations of the VEGFA RNA switch as predicted by SwitchFinder. Middle: SwitchFinder score reflecting the likelihood of a given nucleotide to be involved in two mutually exclusive base pairings. (B) ROC curves of SwitchFinder predictions of RNA switches from the common RFAM families (C) AUC values of RNA switch predictions across the RFAM families for two models: SwitchFinder and SwiSpot (Barsacchi et al. 2016). Each dot represents one RFAM family. The lines show the change in accuracy between the two models. The families that have higher AUC values for SwitchFinder are shown with blue lines; the ones that have higher AUC values for SwiSpot are shown in red. P-value calculated with paired T test.

To discover new families of RNA switches, we aimed to design an approach that does not rely on known sequence motifs, which has been the case for most published software (Wheeler and Eddy 2013; Nawrocki and Eddy 2013; Bengert and Dandekar 2004; Abreu-Goodger and Merino 2005; Chang et al. 2009; Mukherjee and Sengupta 2016). Instead, SwitchFinder uses the sequence to generate an ensemble of secondary structures and their corresponding energy landscape using a Boltzmann equilibrium probability distribution (Ding and Lawrence 2003). It then prioritizes those sequences that show RNA switch-like features, such as having two local minima in close proximity with a relatively small barrier in between (Suppl. Fig. 1B). This approach ensures that RNA switches are described in a generalizable and family-agnostic way. We demonstrated this point by holding individual Rfam families out of the training set and testing whether SwitchFinder would predict the switches correctly (Suppl. Fig. 1C). We observed high performance metrics across all held-out families as measured by the Area Under the Receiver Operating Characteristic curve (AUROC) values (Fig. 1B). We further compared the performance of SwitchFinder to SwiSpot, the state-of-the-art method for family-agnostic riboswitch prediction (Barsacchi et al. 2016), and observed significant improvement of performance across all but one common RNA switch families (Fig. 1C). By relying on biophysical features of the folding energy landscape as opposed to sequence features, SwitchFinder captures a wider variety of RNA switches compared to the existing methods.

In addition to primary sequence, we use RNA secondary structure probing data, when available, to improve SwitchFinder predictions by updating the energy terms of the model. Eukaryotic genomes are large, therefore the models for RNA structural element prediction need to show very high specificity to limit the number of false-positives. Such specificity is difficult to reach by relying on *in silico* RNA folding alone, since RNAs can fold differently *in vivo* and even as a function of cellular state (Beaudoin et al. 2018). However, it is possible to achieve higher specificity if the RNA secondary structure is first probed *in vivo* and the model is then guided by this data. Therefore, we designed SwitchFinder to have an option to update the energy landscape based on RNA secondary structure data (see Methods) and used this functionality to improve our RNA switch predictions iteratively. We named this iterative framework SwitchSeeker. First, we applied the SwitchFinder model using the naive *in silico* folding to the entirety of the 3’ untranslated regions (3’UTR) of the human transcriptome, and chose the 3,750 top candidate switches (of length <= 186 nucleotides) as putative elements. We then probed the secondary structure and function of these candidate RNA switches using two high-throughput *in vivo* screens and fed the resulting structural data into a second iteration of SwitchFinder, which yielded 1,454 high-confidence predictions. Further functional and biochemical validation was carried out *in vivo* (Fig. 2A).

**Fig 2:**
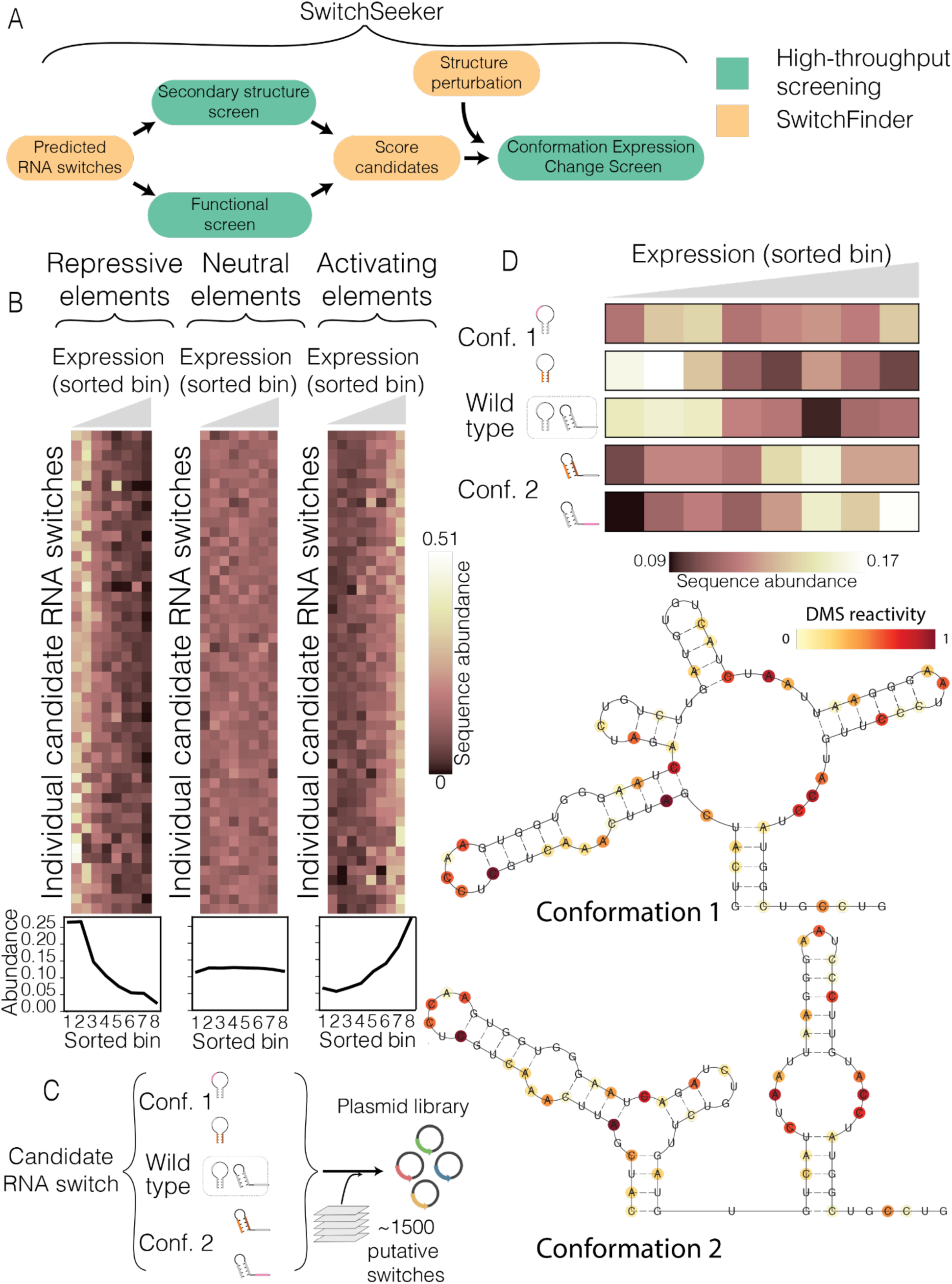
Massively parallel reporter assay captures the functional difference between the conformations of candidate RNA switches. (A) Overview of SwitchSeeker, the platform for RNA switch identification (B) Examples of regulatory elements identified by the functional screen. Each row represents a single candidate RNA switch, each column represents a single bin defined by the reporter gene expression (eGFP fluorescence, normalized by mCherry fluorescence). The value in each cell is the relative abundance of the given RNA switch in the given bin, normalized across the 8 bins. The three plots show examples of candidate switches with repressive, neutral, and activating effects on gene expression. The plots below show cumulative sequence abundances across all the candidate switches within each group. (C) The setup of the massively parallel mutagenesis analysis. For each candidate RNA switch, we design 4 mutated sequence variants. Two of them lock the switch into the conformation 1, and the other two lock it in the conformation 2. A sequence library is then generated (see Suppl. Fig. 2A), where each candidate RNA switch is represented by the 4 mutated sequence variants, along with the wild type sequence. (D) Example of a high-confidence candidate RNA switch identified by the massively parallel mutagenesis analysis. Bottom: two alternative conformations as predicted by SwitchSeeker. The RNA secondary structure probing data collected with the Structure Screen is shown in color. Top: the effect of the candidate RNA switch locked in one or another conformation on reporter gene expression. Each row corresponds to a single sequence variation that locks the RNA switch in one of the two conformations. Each column represents a single bin defined by the reporter gene expression. The value in each cell is the relative abundance of the given RNA switch in the given bin, normalized across the 8 bins.

### Discovery of RNA switches with regulatory function in the human transcriptome

To identify the RNA switches that are both functional and structurally bistable in the cell, we independently performed two high-throughput *in vivo* screens, which we call the “structure screen” and the “functional screen”, respectively. The structure screen differentiates RNAs that exist as an ensemble of two mutually exclusive conformations from those that reside only in a single conformation. The functional screen measures the effect of candidate RNA switches on the expression of a reporter gene. Integrating data from the two screens allowed us to identify the putative RNA switches that are regulatorily active and act as switches *in vivo*. Together with the computational module SwitchFinder, these high-throughput screening strategies comprise the integrated platform SwitchSeeker.

### Large-scale RNA secondary structure probing for improved RNA structural switch predictions

We performed an *in vivo* RNA structure screen using DMS-MaPseq to identify bi-stable RNA structures in the initial pool of 3,750 candidate switches (Zubradt et al. 2017; Mortimer et al. 2012). DMS preferentially modifies unpaired nucleotides resulting in substitutions of adenines and cytosines during the reverse transcription process (Suppl. Fig. 2A). Once the cDNA library is sequenced, the substitution frequency at a given position provides an estimate of nucleotide accessibility, which is typically lower for paired nucleotides versus unpaired. Applying this method to cells expressing the library of candidate RNA switches in a reporter gene context allowed us to collect targeted accessibility measurements with single-nucleotide resolution across all 3,750 candidate RNA switches (Suppl. Fig. 2B,C).

The accessibility of a single nucleotide is a population average of multiple RNA molecules that represent different minima in the RNA folding conformation ensemble. If one conformation dominates within the ensemble, it dominates the DMS-MaPseq reactivity profile; however, if multiple conformations co-exist, they all contribute to the reactivity profile (Morandi et al. 2021; Tomezsko et al. 2020). SwitchSeeker exploits this distinction in nucleotide accessibility to find RNA switches that coexist in a balanced state between two conformations *in vivo*.

### Massively parallel reporter assays identify RNA switches with *in vivo* regulatory functions

We simultaneously sought to explore the potential role of the identified RNA switches in regulating gene expression. We implemented a massively parallel reporter assay (MPRA) (Oikonomou, Goodarzi, and Tavazoie 2014) to functionally interrogate RNA switches *in vivo* (“functional screen”). For this, we tested whether a given RNA switch placed in a 3’ UTR can affect expression of its host mRNA (in this case the eGFP ORF), compared to a control scrambled sequence. We cloned a library of 3,750 candidate RNA switch sequences into a dual eGFP-mCherry fluorescent reporter vector, directly downstream of the eGFP ORF (Suppl. Fig. 2D). We used eGFP fluorescence to measure the effect of candidate RNA switches on gene expression, and we used mCherry fluorescence as an endogenous control. We transduced HEK293 cells with this synthetic library, used flow-cytometry to sort cells by eGFP/mCherry expression ratio, and sequenced the genomic DNA and RNA from the resulting eight pools of cells (Suppl. Fig. 2E, see Methods). Of the candidate RNA switches tested, 536 (14%) showed significant downregulation relative to their scrambled control, and 538 (14%) showed a significant upregulation. We have included representative candidates with repressive, neutral, or activating function in Fig. 2B. While our study focused on characterizing the RNA switches that act in the context of 3’UTRs, the SwitchSeeker framework can be applied to studying other types of RNA switches given appropriate and customized reporter constructs.

To annotate a high-confidence set of RNA switches with regulatory potential in the human transcriptome, we performed a second iteration of SwitchSeeker predictions, guided by the *in vivo* RNA structure probing data. To test the performance of this procedure, we compared the fraction of regulatory active candidate RNA switches among the first and the second iterations of SwitchSeeker prediction, using the functional screen data. We observed a higher fraction of regulatory active RNA switches among the second iteration of SwitchSeeker predictions compared to the first iteration (P = 1e-06, see Suppl. Fig. 2F). This further supports the hypothesis that incorporating the *in vivo* RNA structure probing data improves its performance. We then integrated the high-confidence RNA switches with the massively parallel reporter data. Together these analyses resulted in 1,454 elements that were significant in both screens.

### Massively parallel mutagenesis analysis identifies conformation-specific RNA switch activities

Having identified the candidate RNA switches that affect gene expression, we aimed to assess the degree to which the two stable conformations show divergent regulatory function. For this, we extended our MPRA to include targeted mutations designed to shift the equilibrium between the two conformations of each riboswitch. This additional screen allowed us to identify *bona fide* RNA switches with strong conformation-dependent activity. Starting with the 1,454 high-confidence RNA switches described above, we engineered mutated variants that would lock RNA switches in one of their two predicted conformations. This was achieved by either disrupting or strengthening conformation-specific stem loops. We then performed a massively parallel reporter assay (Suppl. Fig. 2G,H) in which each candidate RNA switch was represented by four additional conformation-specific variants (i.e., two vs. two) (Fig. 2C). We observed a total of 245 RNA switches that differentially regulated reporter gene expression when locked in a specific structural conformation. An example candidate switch (located in the 3’UTR of TCF7) is shown in Fig. 2D; the TCF7 RNA switch landscape has two local minima, corresponding to two alternative conformations supported by *in vivo* DMS-MaPseq data (Fig. 2D, right). Two mutations in different parts of the switch sequence that favor conformation 1 resulted in lower expression of the eGFP reporter. Conversely, two mutations that favor conformation 2 increased eGFP expression. This observation indicates that the two conformations of the TCF7 RNA switch elicit divergent regulatory functions.

### Describing a bi-stable RNA switch in the 3’UTR of RORC

To demonstrate the validity of SwitchSeeker predictions, we sought to characterize the top performing element biochemically. This bistable RNA switch is in the 3’UTR of the RORC mRNA (Fig. 3A); in this RNA switch, the 5’ region can pair either with the 3’ or with the middle region, leading to two mutually exclusive conformations (Fig. 3A). Our measurements indicate that this RNA switch exists in an equilibrium state between the two conformations *in vivo*, and that these two conformations have different effects on the expression of RORC.

**Fig. 3:**
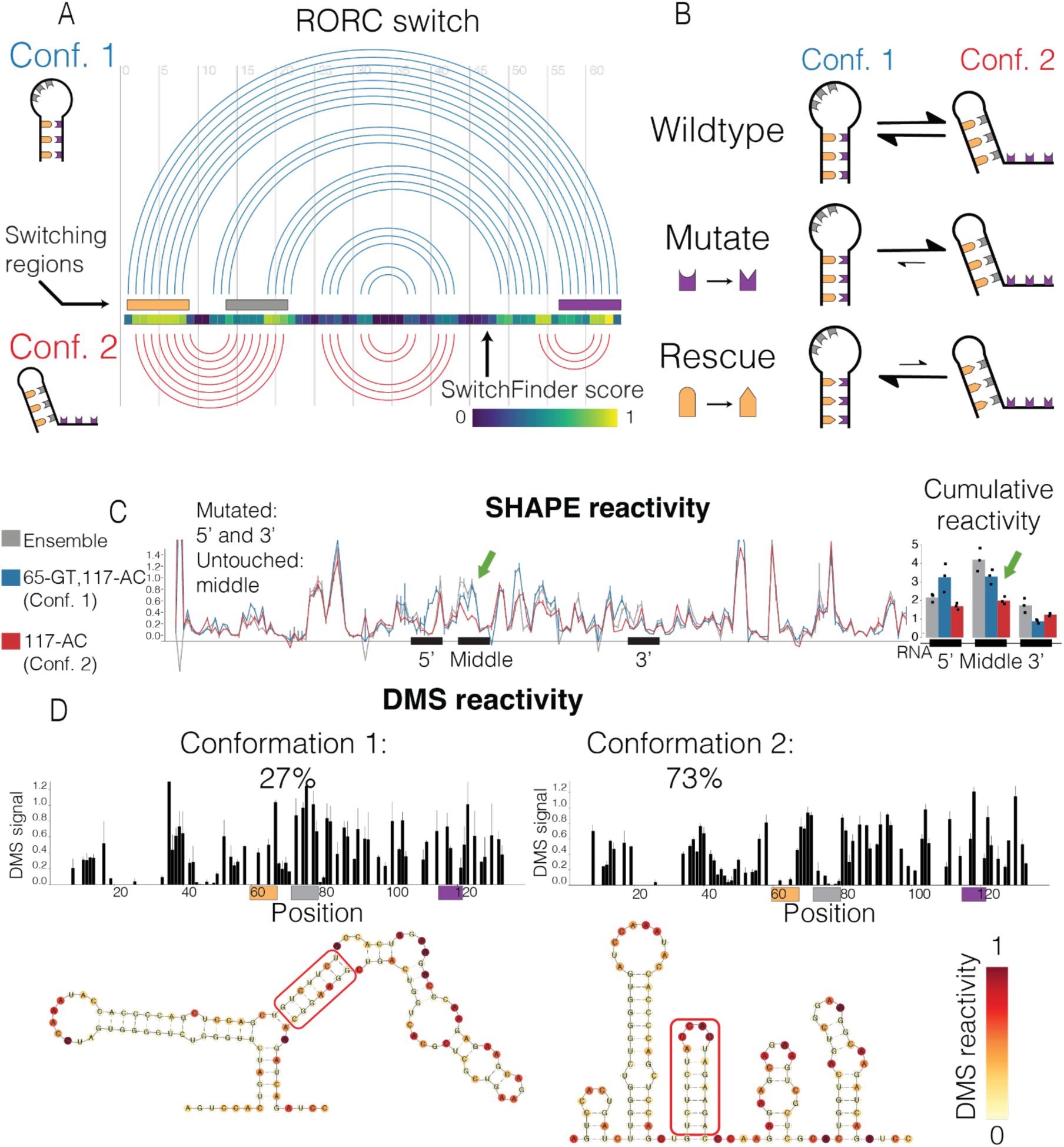
A fragment of RORC 3’UTR forms an ensemble of two alternative structures. (A) Arc representation of the two alternative conformations of the RORC RNA switch as predicted by SwitchSeeker. The two conformations are shown in blue and red, respectively. The predicted switching regions are color coded: the 5’ region is shown in orange, the middle region -in gray, and the 3’ region - in purple. Middle: SwitchSeeker score reflecting the likelihood of a given nucleotide to be involved in two mutually exclusive base pairings. Left: the schematic representations of the two conformations, as used throughout the article. (B) The setup of mutation-rescue experiments. The switching regions are color coded as in (A). A-U and C-G base pairing is shown with compatible shapes (triangle and half-circle). The two conformations of the switch reside in the equilibrium state. Mutation of the 3’ switching region disrupts the base pairing between the 5’ and the 3’ regions. This causes a shift of the equilibrium towards the conformation 2. Rescue mutation of the 5’ switching region restores the base pairing between the 5’ and the 3’ regions, but at the same time it disrupts the base pairing between the 5’ and the middle regions. Therefore, the equilibrium shifts towards conformation 1. (C) In vitro SHAPE reactivity of the RORC RNA switch sequence *in vitro*. Left: SHAPE reactivity profiles for the wild type sequence (in gray) and for the mutation-rescue pair of sequences (blue - “65-GT,117-AC”, red - “117-AC”.). Shown is the average for 3 replicates with the respective error bars. The three switching regions are labeled. The SHAPE reactivity changes in the non-mutated regions are highlighted in bold arrows. Right: barplots of cumulative SHAPE reactivity within the switching regions. The color scheme for the conformations is the same as in the left panel. N replicates = 3. (D) DMS reactivity of the RORC RNA switch *in vivo*. Top: DMS reactivities of the two clusters identified by the DRACO unsupervised deconvolution algorithm (Morandi et al. 2021). The algorithm was run on two replicates independently, and identified the same clusters in both of them. The ratios of the clusters reported by DRACO are 22% to 78% in replicate 1 and 32% to 68% in replicate 2. The ratio shown is an average between the two replicates. The switching regions are shown in color. Bottom: secondary structures of the two conformations of RORC RNA switch predicted by RNAstructure algorithm (Reuter and Mathews 2010) guided by the DMS reactivity data. DMS reactivity is shown in color. The basepairing of 5’ region with either 3’ region (conformation 1) or middle region (conformation 2) are highlighted by a red frame.

To further confirm that the RORC RNA switch exists as an ensemble of two conformations, we performed targeted mutagenesis experiments with *in vitro* RNA SHAPE <u>(Wilkinson et al. 2006)</u> as the read-out. We designed mutation-rescue pairs of sequences that first shift the equilibrium towards one conformation (mutation), and then shift it towards the other conformation (rescue) (Fig. 3B, Suppl. Table 1). We then measured the accessibility of individual nucleotides using the *in vitro* SHAPE assay (Fig. 3C). We observed that mutating the 3’ region (117-AC), which is expected to stabilize conformation 2, reduced the accessibility of the middle region. Conversely, the rescue mutation (65-GT,117-AC) of the 5’ region restored its wild-type accessibility (Fig. 3C). Complementary experiments using the mutation (77-GA) to stabilize conformation 1, and the rescue mutation (63-TC,77-GA) to stabilize conformation 2, had a similar outcome. Even though we did not observe a significant decrease in accessibility of the 3’ region upon the 77-GA mutation, the rescue significantly increased its accessibility (Suppl. Fig. 3A,B). These findings support the role of the three highlighted regions in forming an ensemble of states in which the middle and the 3’ region compete for base pairing to the 5’ region.

To extend our *in vitro* observations to living cells, we performed high-coverage DMS-MaPseq of the RORC switch *in vivo* (Suppl. Fig. 3C). We used a DMS concentration sufficient to cause multiple modifications to the same RNA molecule. This enabled us to cluster reads originating from alternative secondary structures using a state-of-the-art unsupervised computational approach, named DRACO (Morandi et al. 2021). In both biological replicates, DRACO identified two clusters, each representing one of the two conformations, at the approximate ratio of 27% to 73% (Fig. 3D). The profiles of the two clusters did not correlate to each other (p-values of 0.18 and 0.72 in the two replicates 1 and 2), yet the profile of each cluster correlated highly between the two replicates (Suppl. Fig. 3D).

### Single-particle cryo-EM reveals distinct RNA switch tertiary structures

We used single-particle cryo-EM to investigate the tertiary structures of the DRACO clusters. Micrographs of the wild-type RORC RNA switch contain a mixture of compact and extended particles, with features suggestive of RNA secondary structure (Fig 4A). In comparison, particles of the conformation 1 mutant (77-GA) appear mostly compact, while those of the conformation 2 mutant (117-AC) are more highly extended (Fig 4A). Quantitative image processing reveals that wild-type RORC RNA separates into three structural classes labeled A, B, and C, with the Class B structure absent in the (77-GA), and Class A absent in (117-AC) (Fig 4B). These 3D structures demonstrate RNA-like tertiary features, including apparent double-stranded helical segments with a discernible major groove, and typical RNA hairpin elements (Suppl. Fig. 4A). The resolution of these reconstructions is limited to ∼10 Å (Suppl. Fig. 4B), due to the extreme flexibility evinced by the raw micrographs and 2D class averages (Fig 4A, Suppl. Fig. 4C-E), but is sufficient for recognition of RNA fold and handedness. Based on their appearance in the mutant datasets, Class B likely represents conformation 2, while Class A represents conformation 1. Interestingly, Class C is present in all three datasets and may represent a folding intermediate lacking the tertiary interactions made by nucleotides 77 and 117.

**Fig. 4.**
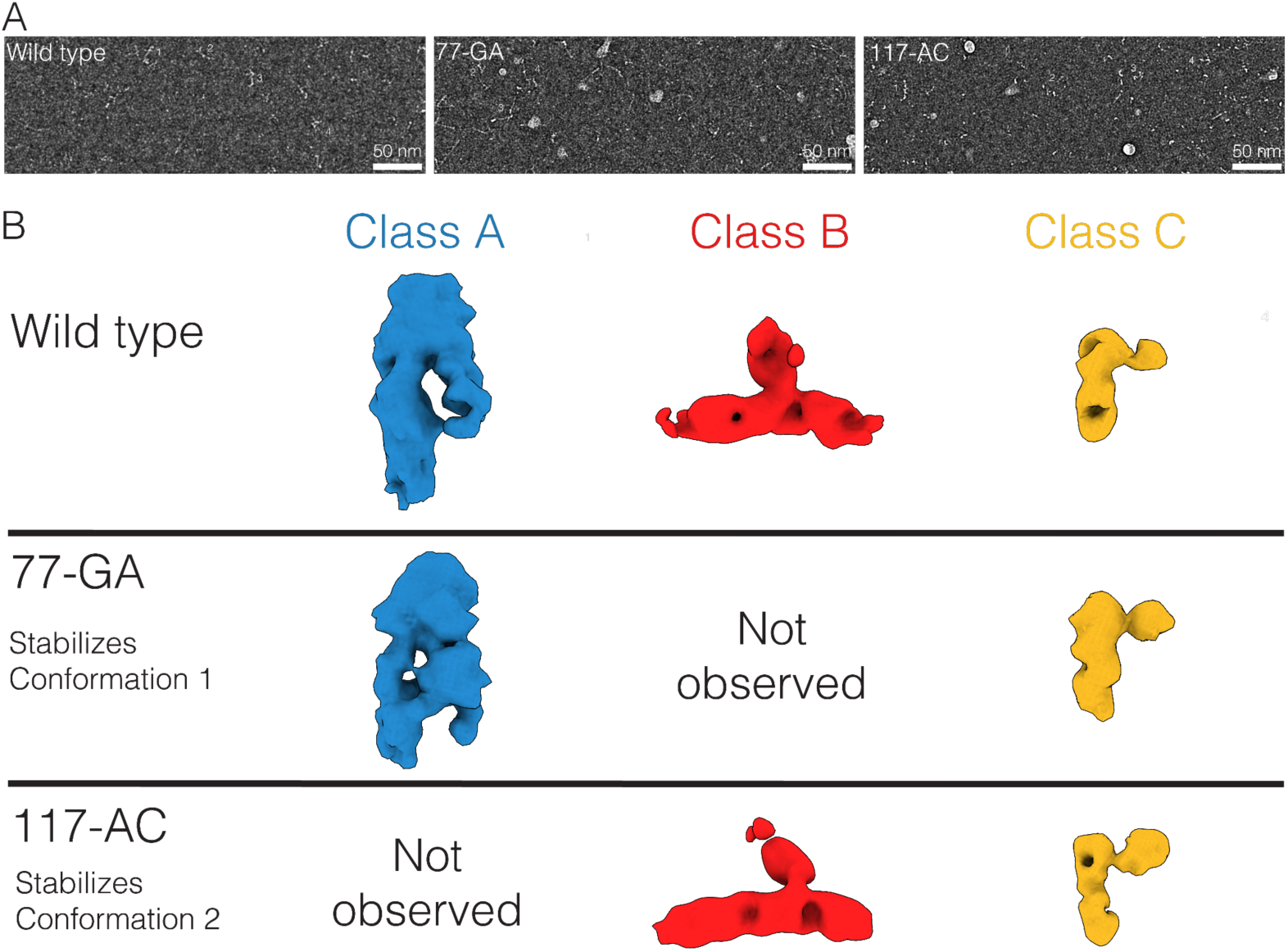
Cryo-EM of RORC 3’ mRNA is consistent with dynamic exchange in a shallow energy landscape. (A) Segments of cryo-EM micrographs of WT RORC mRNA (left), 77-GA mutant (middle), 117-AC mutant (right) show qualitatively different distributions of compact and extended RNA-like particles. Examples with different morphologies are indicated by numeric labels in each panel. Micrographs were phase-flipped, Gaussian filtered, and contrast inverted for display. (B) Cryo-EM of the refolded RORC 3’ mRNA element reveals three classes with RNA-like features (top). Class A is presented in red, Class B in blue, and Class C in yellow. Further cryo-EM imaging and 3D classification of the 77-GA mutant (middle) and 117-AC mutant (bottom) indicate that Class A is present in WT and 77-GA samples but absent from the 117-AC sample, and Class B is conversely present in WT and 117-AC but absent from the 77-GA mutant. Class C is common to all three samples. We thus putatively assign Class A as the conformation 1 state, and Class B as the conformation 2 state. We propose Class 3 to represent a partly folded intermediate which is not disrupted in the mutated constructs.

### The alternative conformations of the *RORC* RNA switch play divergent roles in gene regulation

Having discovered and described the RORC RNA switch as an ensemble of two conformations, we set out to further characterize the divergent regulatory activity of its two states. For this, we generated HEK293 cell lines stably expressing a reporter with the conformation mutant sequences cloned into the 3’UTR of the eGFP coding sequence. We then measured the changes in eGFP expression of each mutant by flow cytometry. We employed two parallel strategies to lock the RNA switch in conformation 1: first, we mutated the middle region so that it cannot pair with the 5’ region, and second, we introduced complementary mutations in both 5’ and 3’ regions, thus preventing base pairing between the 5’ and middle regions. These two orthogonal strategies achieve the same stabilization of secondary structures while modifying different parts of the sequence. Strikingly, both sets of modifications led to similar changes in eGFP expression: locking the RNA switch in conformation 1 increased reporter gene expression compared to wild-type (Fig. 5A). Furthermore, we applied the same two strategies to instead lock the RNA switch in conformation 2 and observed the expected opposite effect: both modifications led to decrease of the reporter gene (Fig. 5A). Therefore, we concluded that the two conformations play divergent functional roles.

**Fig. 5:**
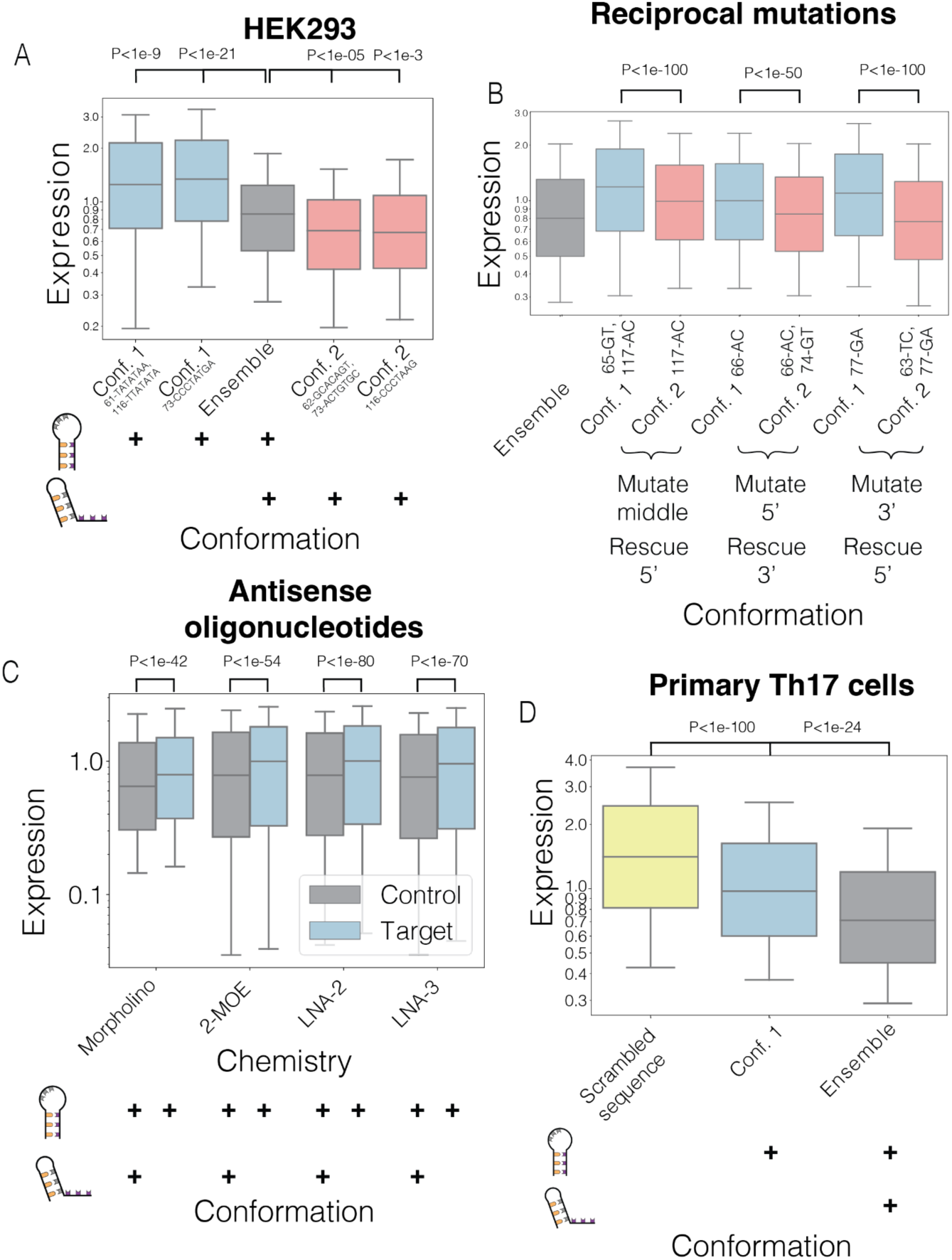
The two alternative conformations of the RORC RNA switch have opposing effects on target gene expression. (A-D) Bar plots of relative expression of the reporter construct. The relative expression values are calculated as the ratios of eGFP and mCherry fluorescence values for individual cells, as measured on a flow cytometer. Confidence intervals for the expression value are shown as calculated from the distribution of N>10.000 individual cells. The dominating conformations of RORC RNA switch are shown in color: conformation 1 - in blue, conformation 2 - in red, wild type - in gray, scrambled sequence with the same dinucleotide content - in yellow. Additionally, diagrams depicting the balance of two conformations in the RNA populations are shown below; the conformations present in the ensemble are highlighted with the “+” sign. P-values were calculated using the Student’s T test. (A) The effect of locking the RORC RNA switch in one or another conformation on reporter gene expression. Bottom: the scheme of the RNA secondary structure ensemble within each sample. The two alternative conformations are depicted as simplified stem-loops; their presence in each sample is shown by a “+” sign. Top: the relative expression of the reporter upon the RORC switch sequence modifications. Each sample is a measurement of a HEK293 cell line with the reporter construct stably integrated in the genome. The mutations left to right: “61-TATATAA,116-TTATATA”, “73-CCCTATGA”, wildtype, “62-GCACAGT,73-ACTGTGC”, “116-CCCTAAG”. (B) The effect of shift in the equilibrium between two conformations of the RORC switch on reporter gene expression. The mutation-rescue experiments were performed as shown in Fig. 3B. Bottom: the direction of mutation-rescue experiments is shown schematically. Top: the relative expression of the reporter upon the RORC switch sequence modifications. The dominating conformation of the switch within a sample is shown in color and labeled under the plot. The mutations left to right: wildtype, “65-GT,117-AC”, “117-AC”, “66-AC”, “66-AC,74-GT”, “77-GA”, “63-TC,77-GA” (C) The effect of antisense oligonucleotides (ASOs) on the reporter expression. A HEK293 cell line with stably integrated reporter construct encoding the RORC RNA switch in the 3’UTR was transfected by ASOs of 4 different chemistries. The targeting ASOs are complementary to the switching region of the RNA switch in the way that would result in a shift towards conformation 1 within the ensemble; the control ASOs have the same nucleotide composition but do not target the RORC RNA switch sequence. Bottom: the RNA switch conformations present in each sample are shown by a “+” sign. Top: the relative expression of the target gene upon the transfection of either targeting or control ASOs. (D) The effect of shift in the equilibrium between two conformations of the RORC switch on reporter gene expression in primary Th17 T cells. Human CD4+ T cells were infected with lentiviral constructs carrying one of the three sequences in the reporter gene’s 3’UTR. The dominating conformations within each sample are shown in color as described above. Bottom: the RNA switch conformations present in each sample are shown by a “+” sign. Top: the relative expression of the target gene upon the transfection of either targeting or control ASOs. The mutations left to right: scrambled RORC RNA switch, “77-GA”, wildtype.

We next tested whether the secondary structure, rather than sequence composition, is the major determinant of the observed modulation in gene expression. To do so, we generated cell lines (as described above) stably expressing the mutant sequences from the rescue-mutation experiments (Fig. 3B) and measured the effect of each mutant on eGFP reporter gene expression. In total, we tested 3 mutation-rescue pairs. In all three cases, we observed lower eGFP expression of the conformation 2 mutant (117-AC), as compared to (77-GA) which favors conformation 1 (Fig. 5B). Taken together, the reciprocal mutation-rescue experiments provide evidence that RNA secondary structure is critical for the conformation-specific function of the RORC RNA switch.

Next, we investigated whether shifting the equilibrium between the two conformations by trans-acting agents, rather than sequence mutations, would have an effect on gene expression. We hypothesized that adding an antisense oligonucleotide (ASO) complementary to a part of the RNA switch sequence could shift the equilibrium between the two conformations to change reporter gene expression. We designed a set of ASOs targeting the middle region of the switch. These ASOs were designed to shift the equilibrium towards conformation 1 and thereby increase eGFP expression. We transfected the stable cell line expressing the RNA switch-containing reporter with ASOs and measured the changes in eGFP expression by flow cytometry. To ensure this effect is not specific to a single ASO design, we varied the oligonucleotide chemistry by using either morpholino, or 2’-O-(2-Methoxyethyl) (2-MOE) oligoribonucleotides, or locked nucleic acids (LNA) as the key modifications. Additionally, we varied the sequence length and the frequency of modifications. In all cases, transfecting cells with an ASO targeting the middle region of the RNA switch resulted in higher eGFP expression compared to a non-targeting ASO of the same chemistry and nucleotide composition (Fig. 5C). Thus, the repressive activity of the RORC RNA switch can be alleviated with the use of trans agents such as ASOs.

The RORC gene encodes nuclear receptor ROR-gamma, and has two protein isoforms that differ by a short N-terminal sequence. The shorter isoform, RORγ, is expressed in many tissues, and is involved in circadian rhythms. The longer isoform, RORγt, is expressed in several subsets of T cells and some lymphoid cells, and is a key driver of Th17 cell type differentiation (Eberl 2017). The RORγt isoform is a drug target for several autoimmune diseases (Zhong and Zhu 2017). Therefore, we also measured the activity of the RORC RNA switch in its endogenous cellular context. We assessed whether the conformation-dependent regulatory function of the RORC RNA switch is observed in Th17 cells. To do this, we infected primary CD4+ cells with lentivirus carrying the reporter and a sequence of interest in the eGFP 3’UTR. The cells were then differentiated into the Th17 cell type (Suppl. Fig. 5, Montoya and Ansel 2017). The presence of the wild type RORC RNA switch strongly decreased eGFP reporter expression compared to a scrambled version of the same sequence (Fig. 5D). On the other hand, the 77-GA mutant decreases the strength of the repression, confirming the activity of the RORC RNA in Th17 cells.

### Genome-scale genetic screens reveal molecular mechanisms underlying the RORC RNA switch

To explore the molecular machinery through which the RORC RNA switch impacts gene expression, we performed genome-wide CRISPRi screens with two distinct eGFP reporters (Suppl. Fig. 6A). The first reporter was designed to identify *trans* factors epistatic to the repressive function of the RORC switch. For this, we created an eGFP construct carrying the wild type RORC RNA switch in its 3’UTR. The second reporter was engineered to assess conformation-dependent activity by inserting the 77-GA RORC switch mutant (favoring conformation 1) in the eGFP 3’UTR. Considering the importance of RORC in T cell biology, we chose the Jurkat T cell leukemia line as the model system for this screen. Jurkat cells were transduced to stably express the dCas9-KRAB CRISPRi protein <u>(Horlbeck et al. 2018)</u>. We infected both reporter cell lines with a genome-wide CRISPRi sgRNA library (Gilbert et al. 2014), sorted cells on a flow cytometer by relative eGFP/mCherry expression, and collected the 25% of cells with highest and lowest eGFP expression (Suppl. Fig. 6A) (de Boer et al. 2020). We hypothesized that knockdown of genes important for the repressive function of the RORC RNA switch would result in higher expression of the reporter gene. Similarly, genes involved in modulating the switch functionality of the two conformations would result in higher reporter expression for the wildtype switch compared to the 77-GA mutant.

To identify factors responsible for the repressive function of RORC RNA switch, we compared the abundance of sgRNAs in the cells with high reporter expression relative to those with lower expression (Fig. 6A). We also performed gene-set enrichment analysis (Korotkevich et al. 2021) to identify the key pathways involved. The most highly enriched pathway was nonsense-mediated decay (NMD)), and several core NMD factors, such as SMG8, UPF1, UPF2, UPF3B were among the highest scoring hits of the screen (Fig. 6A). As expected, among other enriched pathways, many were associated with general gene expression such as translation, ribosome biogenesis, and endoplasmic reticulum stress (Suppl. Fig. 6B). Next, we asked which factors are responsible for the divergent activity of the two conformations. For this, we performed the ratio of ratios test (see methods), by comparing the ratios of sgRNA abundance in low and high expression cells between the two reporter screens: wildtype versus 77-GA mutant (Suppl. Fig. 6C). Interestingly, this comparison also highlighted the NMD pathway as the key contributor (Fig. 6B). Among the highest scoring hits were several core NMD factors; all of them are part of the SURF complex, which is thought to recognize stalled ribosomes when a premature termination codon (PTC) occurs upstream of the exon-junction complex (EJC) (Yamashita 2013). However, the components of EJC did not show any phenotype change in either of the two comparisons (Fig. 6A,B). Therefore, the RORC RNA switch affects gene expression by acting through the EJC-independent NMD pathway, and one conformation of the switch preferentially interacts with this machinery (Yi et al. 2022; Kurosaki, Popp, and Maquat 2019).

**Fig. 6:**
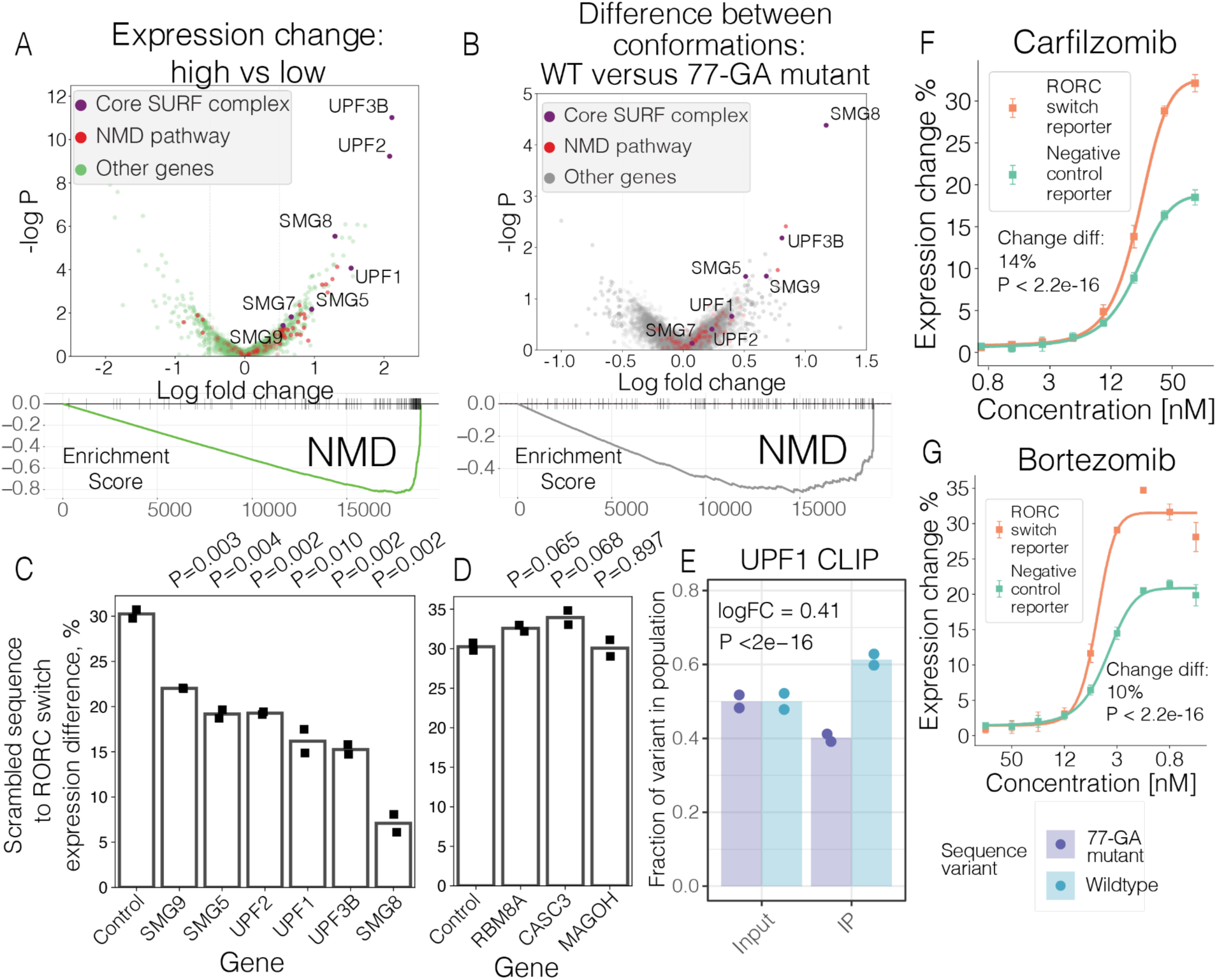
Genome wide CRISPRi screen identifies SURF complex as acting downstream of the RORC RNA switch. (A) The comparison between the high expression and low expression quartiles. Top: volcano plot for gene-wise comparison. The comparison of sgRNA representation between the bottom and the top quantiles by expression (across both cell lines) is represented as a volcano plot. Genes, annotated as part of the nonsense mediated decay (NMD) pathway by GO, are colored in red. The core components of the canonical NMD pathway are colored in purple and labeled. All the other genes are colored in green. Bottom: Gene Set Enrichment Analysis (GSEA) plot for NMD pathway for the above comparison. (B) Comparison of ratios between top and bottom expression quantiles for the two cell lines. Higher values on the X axis indicate that sgRNAs targeting this gene have a stronger effect on reporter gene expression in the WT cell line compared to the RC cell line. Top: “ratio of ratios” comparison (“DESeq2 Testing Ratio of Ratios (RIP-Seq, CLIP-Seq, Ribosomal Profiling)” n.d.) represented as a volcano plot. Genes, annotated as part of the nonsense mediated decay (NMD) pathway by GO, are colored in red. The core components of the canonical NMD pathway are colored in purple and labeled. All the other genes are colored in gray. Bottom: GSEA plot for NMD pathway for the above comparison. (C) (C and D) The effect of knockdown of SURF and EJC complex member proteins on the expression change upon the conformation equilibrium shift. The individual genes were knocked down using the CRISPRi system in both “wild type” and “scrambled” cell lines, then the change of reporter gene expression was measured by flow cytometry (N replicates = 2). The bar plots demonstrate the expression ratios of WT to the scrambled sequence of RORC RNA switch. P-values were calculated using the Student’s T test. (D) The effect of EJC complex knockdown on the expression change upon the conformation equilibrium shift. (E) The fractions of reads carrying the WT RORC switch sequence or 77-GA mutant variant in UPF1 cross-linking and immunoprecipitation (CLIP) library. Left: input RNA libraries, extracted from the WT and 77-GA mutant expressing Jurkat cells, mixed at 1:1 ratio. Right: libraries after anti-UPF1 immunoprecipitation. The fractions are normalized by the variant fractions in the input libraries. (F) (F and G) The effect of proteasome inhibitors bortezomib and carfilzomib on the RNA switch-mediated expression change. K562 cell lines expressing GFP/mCherry double reporter either with or without RORC RNA switch in the 3’UTR of GFP were treated with the drugs for 24 hours, then the reporter gene expression (GFP signal normalized by mCherry signal) was measured by flow cytometry (N replicates = 4). The dose curves for the cell lines expressing or not expressing the WT RORC switch are shown in orange and in green, respectively. (G) The effect of proteasome inhibitor bortezomib on the RNA switch-mediated expression change.

To further confirm this observation, we used CRISPRi to individually knock down the identified core NMD factors in the wild type and 77-GA mutant reporter lines, along with cells expressing a scrambled RORC RNA switch control (“scrambled” cell line). We first asked whether silencing the NMD factors would affect the repressive function of the RNA switch. We compared the expression of eGFP in the wild type and scrambled reporter lines. We observed that silencing the core members of the SURF complex (Fig. 6C), but not of the EJC complex (Fig. 6D), affected the repressive function of the RORC RNA switch sequence on gene expression. Next, we asked whether knockdown of the NMD proteins would reduce the functional difference between the two switch conformations; we compared eGFP expression in the wild type and the 77-GA mutant reporter lines. Consistently, we observed that knocking down the core members of the SURF complex, and not of the EJC complex (Suppl. Fig. 6D), reduced this difference. This data demonstrates that NMD acts preferentially on the Conformation 2 of the RORC RNA switch.

Given that UPF1 can bind highly structured RNA regions (Fischer et al. 2020), we hypothesized that UPF1 might bind the two RORC RNA switch conformations with different affinities. To test this hypothesis, we mixed together the wild type and the 77-GA mutant reporter lines at a 1:1 ratio, and measured the difference in UPF1 affinity by targeted crosslinking immunoprecipitation (CLIP) followed by targeted sequencing. We observed that the wildtype RORC UTR sequence was significantly more abundant among the UPF1 CLIP tags than its 77-GA mutant (Fig. 6E). To further confirm the conformation-specific interaction of the RORC switch with UPF1, we mixed together cell lines expressing two mutants that lock the RNA switch in the opposite conformations: 77-GA (locks the conformation 1) and 116-CCCTAAG (locks the conformation 2), and measured the UPF1 binding differences as described above. As expected, we observed higher fraction of reads aligning to the 116-CCCTAAG mutant sequence than to the 77-GA mutant (Suppl. Fig. 6E). The observed difference in variant sequence fractions was larger for the mix of two opposite mutants than for the mix of 77-GA mutant with the wildtype sequence (LogFC values of 1.12 vs 0.41). Taken together, our results indicate differential UPF1 binding to the two conformations of the RNA switch.

Since the canonical NMD pathway causes the proteins translated from aberrant mRNA to be degraded by the proteasome (Kuroha, Tatematsu, and Inada 2009), we expect the RORC RNA switch to target its gene product for degradation by the proteasome. To confirm this, we treated two dual reporter cell lines with the proteasome inhibitor carfilzomib, one line expressing the RORC RNA switch (the “wild type” line), and the other expressing the scrambled control. As expected, proteasome inhibition resulted in a significantly larger change in eGFP expression in the “wild-type” line relative to the “scrambled” cell line (Fig. 6F). To verify that the observed effect is due to proteasome inhibition rather than a side effect of carfilzomib, we treated cells with bortezomib, another proteasome inhibitor which acts through a different mechanism, and observed the same effect (Fig. 6G). Therefore, our data suggests that proteasomal degradation is involved in the repressive activity of the RORC RNA switch on gene expression.

## Discussion

In the past, RNA switches were discovered through either comparative genomics analysis or biochemical experimentation. Comparative genomic analysis searches for conserved positions within non-coding RNA regions and has proven effective for identifying cis-regulatory elements in bacteria, including riboswitches and transcription factor binding sites (Rodionov 2007). The biochemical approach involves measuring the affinity of a putative RNA switch to its ligands and analyzing the conformational change caused by the binding event. Both approaches were used to discover the first known RNA switches in bacteria (Epshtein, Mironov, and Nudler 2003; Winkler, Nahvi, and Breaker 2002). However, no specific algorithms have been created to identify RNA switches in eukaryotic transcriptomes. In Eukaryotes, mRNA secondary structure is highly dynamic; multiple studies have shown that RNA structure vastly differs when measured *in vitro* vs *in vivo* (Rouskin et al. 2014), and that multiple cellular processes can rearrange mRNA secondary structures (Sun et al. 2019). Although the functional importance of individual RNA structure rearrangements has been studied, such as RNA thermosensors (Shamovsky et al. 2006), the extent to which structural switches control gene expression in eukaryotes remains largely unexplored. There are several reasons why models trained on bacteria cannot be easily applied to metazoans, including the larger sequence search space in eukaryotic transcriptomes, which hinders the application of pre-trained models due to high false-positive counts (Ureta-Vidal, Ettwiller, and Birney 2003), and poor sequence conservation of many eukaryotic RNA regulatory elements, which limits the applicability of comparative genomics analyses (Backofen et al. 2018; Leypold and Speicher 2021). However, the SwitchSeeker methodology provides a comprehensive, integrative platform for studying RNA switches in eukaryotic transcriptomes. It covers all the steps of discovery, from *de novo* predictions to uncovering the mechanism of individual RNA switches, and overcomes the limitations of exisiting methodologies by integrating the toolkits of biochemistry, systems biology, and functional genomics. With its scalability to complete transcriptomes, SwitchSeeker offers a way towards comprehensive characterization of RNA switches across the tree of life.

Recent advances in genomic technologies were a key contributor in our ability to carry out this systematic search for RNA switches. RNA secondary structure probing techniques, such as DMS-seq and SHAPE-seq, have enabled researchers to study multiple alternative conformations of RNA molecules (Tomezsko et al. 2020; Morandi et al. 2021). Additionally, recent advancements in single-particle cryo-EM and computational modeling have enabled the determination of the 3D folds of some RNA molecules (Kappel et al. 2020), despite their small size and intrinsic flexibility. This has opened up opportunities to study the functional differences between alternative RNA conformations and their role in gene expression control. Our DMS-MapSeq and cryo-EM data suggest that the RORC 3’ mRNA element inhabits a shallow energy landscape with two rugged minima linked to two major molecular conformations (Fig. 7A). Mutations that stabilize one conformation or another alter the energy landscape and the distribution of states. Experimental structure determination thus validates the SwitchSeeker approach to identification of RNA molecules with these types of bistable energy landscapes. The RORC RNA switch was functionally characterized using CRISPRi, which revealed that its control of gene expression is mediated by the SURF complex of the EJC-independent NMD. We propose that UPF1 preferentially recognizes switch conformation 2 over conformation 1, and that the recruitment of the SURF complex by UPF1 in turn leads to decreased gene expression through proteasome-mediated degradation of any translation products, and mRNA decay preventing repeated rounds of translation (Fig. 7B). Point mutants that modulate the conformational equilibrium likewise cause small but significant changes in the degree of gene repression by the switch. Switch sequence variation can likely provide nearly continuous modulation of SURF recruitment and NMD activation. However, questions remain regarding the pathways that trigger the switching between the two conformations.

**Fig. 7:**
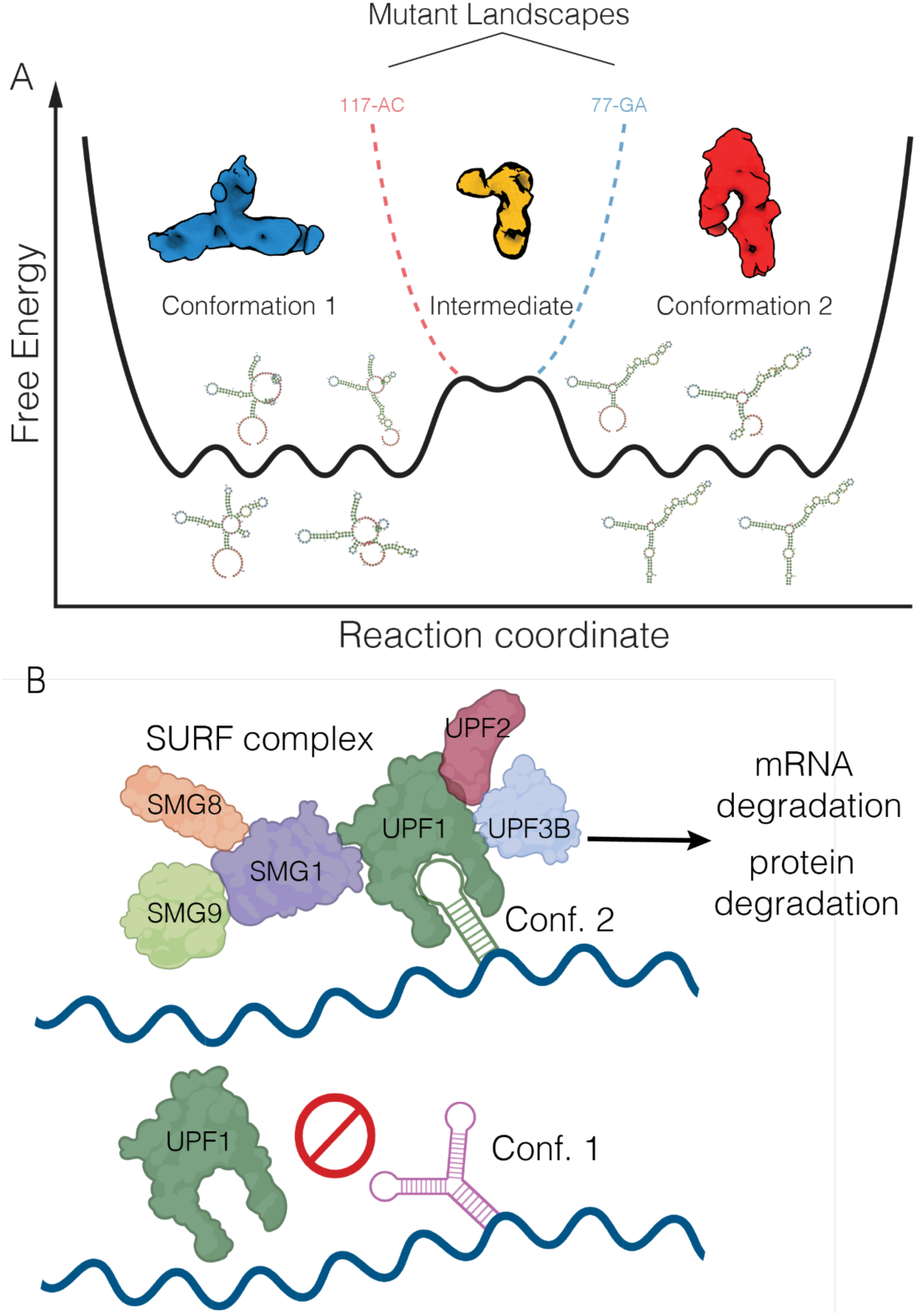
The proposed mechanism of RORC RNA switch functioning. (A) Schematic for a shallow energy landscape of the RORC 3’ mRNA element. Shallow global minima characterizing the conformation 1 (cryo-EM Class A) and conformation 2 (cryo-EM Class B) structures themselves comprise multiple local minima in which various secondary structure elements fold or unfold while preserving overall tertiary structure and biological activity. These local minima are illustrated by secondary structure models for various DRACO cluster members. The two global minima are separated by a kinetic barrier that represents a partially folded intermediate (cryo-EM Class 3). The two dashed lines indicate alterations to the global landscape exhibited by the mutant sequences, red for the 77-GA mutant and blue for the 117-AC mutant. These altered landscapes eliminate one of the global minima without disrupting the intermediate. (B) Proposed mechanism of RORC RNA switch. The RNA switch exists in an ensemble of two states. One of them is recognized by the SURF complex; such recognition triggers mRNA degradation (likely mediated by SMG5) and protein degradation (mediated by proteasome), thus affecting gene expression.

We have observed that a large number of regulatory elements in the human transcriptome act as RNA switches. This implies that conformation-dependent modes of gene expression control are likely widespread in the human transcriptome. The regulatory information contained in dynamic RNA structural elements is an under-explored area of gene expression control that plays a fundamental role in health and disease. Understanding the regulatory grammar of RNA switches throughout the transcriptome is a crucial step towards a more comprehensive comprehension of post-transcriptional gene expression control. In our study, we chose stringent criteria for selecting RNA switches, requiring them to be bistable *in vivo*, meaning they populate two mutually exclusive structural conformations. However, this criterion may not be applicable to all functional RNA switches, which may be bistable only under specific conditions or in specific cell types. For instance, it has been demonstrated recently that HIV-1 TAR RNA forms a rare and short-lived yet biologically active conformation (Kelly et al. 2022). Additionally, RNA switches may be multistable and provide even more complex regulation. Developing further methodologies is necessary to capture these attenuated regulatory elements.

RNA switches function through different mechanisms. The known examples of human RNA switch mechanisms include mutually exclusive binding of RBPs by two different conformations (Ray et al. 2009) and m6A modification-based switching (Liu et al. 2015). In this study, we presented a novel RNA switch that operates via the NMD pathway, and we believe that other RNA switches may also exploit various RNA metabolic pathways. For example, we have recently shown that specific RNA structures cause aberrant splicing in metastatic cancers through binding SNRPA1 (Fish et al. 2021). Thus, future studies will likely find a wider variety of RNA switches than those discovered in the present study, under steady-state conditions. Importantly, SwitchSeeker is general with respect to the mechanism: it can identify general RNA structural switches and reveal their mechanisms. We anticipate that many RNA switches will be employed as therapeutic handles for treating genetic diseases: in this study, we demonstrated that it is possible to control gene expression by shifting the balance between the two RNA conformations with therapeutics such as ASOs. Understanding the regulatory programs that govern RNA secondary structure switching will provide a mechanistic understanding of gene expression control, which will be invaluable for studying RNA switches in various contexts such as development and disease. SwitchSeeker is available for use and adaptation, and we hope it will pave the way for further discoveries in RNA-based regulation in eukaryotes.

## Methods

### SwitchFinder: detailed description of the algorithm

#### Conflicting basepairs identification

Conflicting basepairs were detected using a modification of MIBP algorithm developed by L. Lin and W. McKerrow (Lin et al. 2018). First, a large number of folds (default N=1000) is sampled from the Boltzmann distribution. If structure probing data (such as DMSseq or SHAPEseq) is provided, the Boltzmann distribution modeling software (part of RNAstructure package (Reuter and Mathews 2010)) incorporates the data as a pseudofree energy change term. Then, the basepairs are filtered: the basepairs that are present in almost all the folds or are absent from almost all the fold are removed from the further analysis. Then, mutual information for each pair of basepairs is estimated. To do so, each basepair is represented as a binary vector of length N, where N is the number of folds considered; in this binary vector, a given fold is represented as 1 if this basepair is present there, or as 0 if it’s not. Mutual information between each two basepairs is calculated as in (Cover and Thomas 2006). This results in a M by M table of mutual information values, where M is the number of basepairs considered. Then, the sum of each row of the square table is calculated. In the resulting vector K of length M, each basepair is represented by a sum of mutual information values of across all the other basepair. Then, only the basepairs whose sum of mutual information values passes the threshold of U * MAX(K) is considered, where U is a parameter (default value 0.5). We call the basepairs that pass this threshold the “conflicting basepairs”.

#### Conflicting stems identifications

Once the conflicting basepairs were identified, they get assembled into conflicting stems, or series of conflicting basepairs that directly follow each other and therefore could potentially form a stem-like RNA structure. More specifically, the basepairs (a, b) and (c, d) form a stem if either (a == c - 1) & (b == d +1), or (a == c + 1) & (b == d - 1). The stem is defined as a pair of intervals ((u, v), (x, y)), where v - u == y - x. Then, the conflicting stems get filtered by length: only the stems that are longer than a certain threshold value (default value: 3) are considered. Among these stems, the stems that directly conflict with each other are identified. Two stems ((u_1_, v_1_), (x_1_, y_1_)) and ((u_2_, v_2_), (x_2_, y_2_)) conflict with each other if there is an overlap longer than a threshold value between either (u_1_, v_1_) and (u_2_, v_2_), or (u_1_, v_1_) and (x_2_, y_2_), or (x_1_, y_1_) and (u_2_, v_2_), or (x_1_, y_1_) and (x_2_, y_2_). Default threshold value is 3. The pairs of conflicting stems are sorted by the average value of their K values (sums of mutual information). The highest scoring pair of conflicting stems is considered the winning prediction, representing the major switch between two of the local minima present in the energy folding landscape of the given sequence. If not pairs of conflicting stems pass the threshold, SwitchFinder reports that no potential switch is identified for the given sequence.

#### Identifying the two conflicting structures

Given the prediction of the two conflicting stems, the folds that represent the two local minima of the energy folding landscape are predicted. Importantly, SwitchFinder focuses on optimizing the prediction accuracy, as opposed to the commonly used approach of energy minimization (Lu, Gloor, and Mathews 2009). MaxExpect program from the RNAstructure package (Reuter and Mathews 2010) is used; the basepairings of each of the conflicting stems are provided as folding constraints (in Connectivity Table format). The two predicted structures are further referred to as conformations 1 and 2.

Activation barrier estimation. The RNApathfinder software (Dotu et al. 2010) is used to estimate the activation energy needed for a transition between the conformations 1 and 2.

#### Classifier for prediction of RNA switches

The curated representative alignments for each of the 50 known riboswitch families were downloaded from RFAM database (Kalvari et al. 2021). Each sequence gets complemented by its shuffled counterpart (while preserving dinucleotide frequencies, see (*altschulEriksonDinuclShuffle.py at Master · wassermanlab/BiasAway* n.d.)). For all the sequences, the two conflicting conformations, their folding energies and their activation energies were predicted as above. To estimate the performance of SwitchFinder for a given riboswitch family, all the sequences from this family are placed into the test set, while all the sequences from the other families are placed into the training set. Then, linear regression model is trained on the training set, where the response variable is binary and indicates whether the sequence is a real riboswitch or is a shuffled counterpart, and the predictor variables are the average folding energy of the two conformations and the activation energy of the transition between them. The trained linear regression model is then run on the test set, and its performance is estimated with the receiver operating characteristic curve.

#### Prediction of RNA switches in human transcriptome

The coordinates of 3’UTRs of the human transcriptome were downloaded from UCSC Table Browser (Karolchik, Hinrichs, and Kent 2012), table tb_wgEncodeGencodeBasicV28lift37. The sequences of 3’UTRs were cut into overlapping fragments of 186 nt in length (with the overlaps of 93nt). For all the sequences, the two conflicting conformations, their folding energies and their activation energies were predicted as above. A linear regression model was trained as described above on all the 50 known riboswitch families. The model was applied to the 3’UTR fragments from the human genome, and the fragments were sorted according to the model prediction scores. The top 3750 predictions were selected for further investigation.

#### Mutation generation

In order to shift the RNA conformation ensemble towards one or another state, mutations of two types were introduced.

(1) “Strengthen a stem” mutations: given two conflicting stems ((u_1_, v_1_), (x_1_, y_1_)) and ((u_2_, v_2_), (x_2_, y_2_)), one of the stems (for example, the first one) was changed in a way that would preserve its basepairing but deny the possibility of forming the second stem. To do so, the nucleotides in the interval (u_1_, v_1_) were replaced with all possible sequences of equal length, and the nucleotides (x_1_, y_1_) were replaced with the reverse complement sequence. Then, the newly generated sequences were filtered by two predetermined criteria: (i) the second stem can not form more than a fraction of its original base pairs (default value 0.6), (ii) the modified first stem can’t form long paired stems with any region of the existing sequence (default threshold length 4). The sequences that passed both criteria were ranked by the introduced change in the sequence nucleotide composition; the mutations that changed the nucleotide composition the least were chosen for further analysis. Each mutated sequence was additionally analysed by SwitchFinder to ensure that the Boltzmann distribution is heavily shifted towards the desired conformation.

(2) “Weaken a stem” mutations: given two conflicting stems ((u_1_, v_1_), (x_1_, y_1_)) and ((u_2_, v_2_), (x_2_, y_2_)), one of the stems (for example, the second one) was changed in a way this stem wouldn’t be able to form anymore, while the basepairing of the other stem (in this example, the first stem) would be preserved. To do so, the nucleotides in either of the intervals (u_2_, v_2_) or (x_2_, y_2_) were replaced with all possible sequences of equal length. The newly generated sequences were filtered by three predetermined criteria: (i) the first stem stays unchanged, (ii) the second stem can not form more than a fraction of its original base pairs (default value 0.6), (ii) the modified part of the sequence can’t form long paired stems with any region of the existing sequence (default threshold length 4). The sequences that passed all the criteria were ranked by the introduced change in the sequence nucleotide composition; the mutations that changed the nucleotide composition the least were chosen for further analysis. Each mutated sequence was additionally analysed by SwitchFinder to ensure that the Boltzmann distribution is heavily shifted towards the desired conformation.

#### Code availability

The source code of SwitchFinder is available at https://github.com/goodarzilab/SwitchFinder.

### Cell culture

All cells were cultured in a 37°C 5% CO2 humidified incubator. The HEK293 cells (ATCC CRL-3216) were cultured in DMEM high-glucose medium supplemented with 10% FBS, L-glutamine (4 mM), sodium pyruvate (1 mM), penicillin (100 units/mL), streptomycin (100 μg/mL) and amphotericin B (1 μg/mL) (Gibco). The Jurkat cell line was cultured in RPMI-1640 medium supplemented with 10% FBS, glucose (2 g/L), L-glutamine (2 mM), 25 mM HEPES, penicillin (100 units/mL), streptomycin (100 μg/mL) and amphotericin B (1 μg/mL) (Gibco). All cell lines were routinely screened for mycoplasma with a PCR-based assay.

### Cryo-EM Sample Preparation and Data Collection

3.5 µl of target mRNA at an approximate concentration of 1.5 mg/mL was applied to gold, 300 mesh TEM grids with a holey carbon substrate of 1.2/1.3 µm spacing (Quantifoil). The grids were blotted with #4 filter papers (Whatman) and plunge frozen in liquid ethane using a Mark IV Vitrobot (Thermo Fisher), with blot times of 4-6 s, blot force of -2, at a temperature of 8°C and 100% humidity. All grids were glow discharged in an easiGlo (Pelco) with rarefied air for 30 s at 15 mA, no more than 1 hour prior to preparation. Duplicate WT and mutant RNA specimens were imaged under different conditions on several microscopes as per Table S8; all were equipped with K3 direct electron detector (DED) cameras (Gatan), and all data collection was performed using SerialEM (Mastronarde 2003). Detailed data collection parameters are listed in Table S8.

### Cryo-EM Image processing

Dose-weighted and motion-corrected sums were generated from raw DED movies on-the-fly during data collection using UCSF MotionCor2 (Zheng et al. 2017). Images from super-resolution datasets were downsampled to the physical pixel size before further processing. CTF estimation was performed in CTFFIND4 (“Ctffind4” n.d.), followed by neural-net based particle picking in EMAN2 (Tang et al. 2007). 2D classification, *ab initio* 3D classification, and gold-standard refinement were done in cryoSPARC (Punjani et al. 2017). CTFs were then re-estimated in cryoSPARC and particles repicked using low-resolution (20 Å) templates generated from chosen 3D classes. Extended datasets were pooled where appropriate, and particle processing was repeated through gold-standard refinement as before. All structure figures were created using UCSF ChimeraX (Goddard et al. 2018). Further details are given in Table S7 and Extended Data Figure 4.

### Reporter vector design and library cloning

First, mCherry-P2A-Puro fusion was cloned into the BTV backbone (Addgene #84771). Then, the vector was digested with MluI-HF and PacI restriction enzymes (NEB), with the addition of rSAP (NEB). The digested vector was purified with Zymo DNA Clean and Concentrator-5 kit.

DNA oligonucleotide libraries (one for functional screen and one for massively parallel mutagenesis analysis) consisting of 7500 sequences total were synthesized by Agilent. The second strand was synthesized using Klenow Fragment (3’ → 5’ exo-) (NEB). The dsDNA library was digested with Mlul-HF and Pad restriction enzymes (NEB) and run on a 6% TBE polyacrylamide gel. The band of the corresponding size was cut out and the gel was dissolved in the DNA extraction buffer (10 mM Tris pH 8, 300 mM NaCI, 1 mM EDTA). The DNA was precipitated with isopropanol. The digested DNA library and the digested vector were ligated with T4 DNA ligase (NEB). The ligation reaction was precipitated with isopropanol and transformed into MegaX DH10B T1R Electrocompetent Cells (Thermo Fisher). The library was purified with ZymoPURE II Plasmid Maxiprep Kit (Zymo). The representation of individual sequences in the library was verified by sequencing the resulting library on MiSeq instrument (lllumina).

### Massively Parallel Reporter Assay

The DNA library was co-transfected with pCMV-dR8.91 and pMD2.G plasmids using TransIT-Lenti (Mirus) into HEK293 cells, following the manufacturer’s protocol. Virus was harvested 48 hours post-transfection and passed through a 0.45 µm filter. HEK293 cells were then transduced overnight with the filtered virus in the presence of 8 µg/mL polybrene (Millipore); the amount of virus used was optimized to ensure the infection rate of ∼20%, as determined by flow cytometry The infected cells were selected with 2 µg/mL puromycin (Gibco). Cells were harvested at 90%–95% confluency for sorting and analysis on a BD FACSaria II sorter. The distribution of mCherry to GFP ratios was calculated. For sorting a library into subpopulations, we gated the population into 8 bins each containing 12.5% of the total number of cells. A total of 1.2 million cells were collected for each bin to ensure sufficient representation of sequence in the population in two replicates each. For each subpopulation, we extracted gDNA and total RNA with the Quick-DNA/RNA Miniprep kit. gDNA was amplified by PCR with Phusion polymerase (NEB) using primers CAAGCAGAAGACGGCATACGAGAT -i7 - GTGACTGGAGTTCAGACGTGTGCTCTTCCGATCACTGCTAGCTAGATGACTAAACGCG and AATGATACGGCGACCACCGAGATCTACAC - i5 - ACACTCTTTCCCTACACGACGCTCTTCCGATCTGTGGTCTGGATCCACCGGTCC. Different i7 indices were used for 8 different bins, and different i5 indices were used for the two replicates. RNA was reverse transcribed with Maxima H Minus Reverse Transcriptase (Thermo Fisher) using primer CTCTTTCCCTACACGACGCTCTTCCGATCTNNNNNNNNNNNTGGTCTGGATCCACCGGTCCGG. The cDNA was amplified with Q5 polymerase (NEB) using primers CAAGCAGAAGACGGCATACGAGAT - i7 - GTGACTGGAGTTCAGACGTGTGCTCTTCCGATCCTGCTAGCTAGATGACTAAACGC and CAAGCAGAAGACGGCATACGAGAT - i5 - GTGACTGGAGTTCAGACGTGTGCTCTTCCGATCTTACCCGTCATTGGCTGTCCA. Different i7 indices were used for 8 different bins, and different i5 indices were used for the two replicates. The amplified DNA libraries were size purified with the Select-a-Size DNA Clean & Concentrator MagBead Kit (Zymo). Deep sequencing was performed using the HiSeq4000 platform (Illumina) at the UCSF Center for Advanced Technologies.

The adapter sequences were removed using cutadapt (Martin 2011). For RNA libraries, the UMI was then removed from the reads and appended to read names using UMI tools (Smith, Heger, and Sudbery 2017). The reads were matched to the fragments using bwa mem command. The reads were counted using featureCounts (Liao, Smyth, and Shi 2014). The read counts were normalized using median of ratios normalization (Anders and Huber 2010). One-way chi-square test statistic was used to estimate how different its distribution across the sorting bins is from the null hypothesis (i.e. uniform distribution). mRNA stability was estimated by comparing the RNA and DNA read counts with MPRAnalyze (Ashuach et al. 2019).

### DMS-MaPseq

DMS-MaPseq was performed as described in (Fish et al. 2021). Briefly, HEK293 cells were incubated in culture with 1.5% DMS (Sigma) at room temperature for 7 minutes, the media was removed, and DMS was quenched with 30% BME. Total RNA from DMS-treated cells and untreated cells was then isolated using Trizol (Invitrogen). RNA was reverse transcribed using TGIRT-III reverse transcriptase (InGex) and target-specific primers. PCR was then performed to amplify the desired sequences and to add Illumina compatible adapters. The libraries were then sequenced on a MiSeq instrument using MiSeq micro kit v2, 300 cycles (Illumina).

Pear (v0.9.6) was used to merge the paired reads into a single combined read. The UMI was then removed from the reads and appended to read names using UMI tools (v1.0). The reads were then reverse complemented (fastx toolkit) and mapped to the amplicon sequences using bwa mem (v0.7). The resulting bam files were then sorted and deduplicated (umi_tools, with method flag set to unique). The alignments were then parsed for mutations (CTK). The mutation frequency at every position was then reported. The signal normalization was performed using boxplot normalization as in (Low and Weeks 2010). The top 10% of positions with the highest mutation rates were considered outliers as suggested in (Hajdin et al. 2013). The clustering of DMS-MaPseq signal was performed with DRACO (Morandi et al. 2021).

### SHAPE chemical probing of RNAs

Chemical probing and mutate-and-map experiments were carried out as described previously (Palka et al. 2020). Briefly, 1.2 pmol of RNA was denaturated at 95°C in 50 mM Na-HEPES, pH 8.0, for 3 min, and folded by cooling to room temperature over 20 min, and adding MgCl2 to 10 mM concentration. RNA was aliquoted in 15 µL volumes into a 96-well plate and mixed with nuclease-free H2O (control), or chemically modified in the presence of 5 mM 1-methyl-7-nitroisatoic anhydride (1M7) (Turner, Shefer, and Ares 2013), for 10 min at room temperature. Chemical modification was stopped by adding 9.75 µL quench and purification mix (1.53 M NaCl, 1.5 µL washed oligo-dT beads, Ambion), 6.4 nM FAM-labeled, reverse-transcriptase primer (/56- FAM/AAAAAAAAAAAAAAAAAAAAGTTGTTCTTGTTGTTTCTTT), and 2.55 M Na-MES. RNA in each well was purified by bead immobilization on a magnetic rack and two washes with 100 µL 70% ethanol. RNA was then resuspended in 2.5 µL nuclease-free water prior to reverse transcription.

RNA was reverse-transcribed from annealed fluorescent primer in a reaction containing 1× First Strand Buffer (Thermo Fisher), 5 mM DTT, 0.8 mM dNTP mix, and 20 U of SuperScript III Reverse Transcriptase (Thermo Fisher) at 48°C for 30 min. RNA was hydrolyzed in the presence of 200 mM NaOH at 95°C for 3 min, then placed on ice for 3 min and quenched with 1 volume 5 M NaCl, 1 volume 2 M HCl, and 1 volume 3 M sodium acetate. cDNA was purified on magnetic beads, then eluted by incubation for 20 min in 11 µL Formamide-ROX350 mix (1000 µL Hi-Di Formamide (Thermo Fisher) and 8 µL ROX350 ladder (Thermo Fisher). Samples were then transferred to a 96-well plate in “concentrated” (4 µL sample + 11 µL ROX mix) and “dilute” (1 µL sample + 14 µL ROX mix) for saturation correction in downstream analysis. Sample plates were sent to Elim Biopharmaceuticals for analysis by capillary electrophoresis.

### ASO infection

Antisense oligonucleotides were purchased from Integrated DNA Technologies; the Morpholino ASOs were purchased from Gene Tools LLC (see sequences in Data file S9). 95.000 HEK293 cells were seeded into a well of a 24-well cell culture treated plate in a total volume of 500 ul. 24 hours later, either 1 nmol of Morpholino ASO together with 3 ul of EndoPorter reagent (Gene Tools LLC), or 6 pmol of other ASO were added to each well. 48 hours later, the mCherry and eGFP fluorescence was measured on BD FACSCelesta Cell Analyzer.

### CRISPRi screen

Reporter screens were conducted using established flow cytometry screen protocols (Gilbert et al. 2014)( Horlbeck et al., 2016; Sidrauski et al., 2015). Jurkat cells with previously verified CRISPRi activity were used (Horlbeck et al., 2018). The CRISPRi-v2 (5 sgRNA/TSS, Addgene: Cat#83969) sgRNA library was transduced into Jurkat cells at an MOI < 0.3 (BFP+ cell percentages were ∼30%). For the flow based CRISPRi screen with the Jurkat cells, the sgRNA library virus was transfected at an average of 500x coverage after transduction (Day 0). Puromycin (1 µg/mL) selection for positively-transduced cells was performed 48 hours (Day 2) and 72 hours (Day 3) post transduction (Day 3). On Day 11, cells were collected in PBS and sorted with the BD FACSAria™ Fusion cell sorter. Cells were gated into the 25% of cells with the highest ratio between GFP and mCherry fluorescence intensity and 25% of cells with the lowest ratio. The screens were performed with two conditions: cells with a wildtype RORC Riboswitch-GFP reporter and a mutated Riboswitch reporter. Screens were additionally performed in duplicate. After sorting, genomic DNA was harvested (Macherey-Nagel Midi Prep kit) and amplified using NEB Next Ultra II Q5 master mix and primers containing TruSeq Indexes for NGS analysis. Sample libraries were prepared and sequenced on a HiSeq 4000. Guides were then quantified with the published ScreenProcessing (https://github.com/mhorlbeck/ScreenProcessing) method and phenotypes generated with an in-house processing pipeline, iAnalyzer (https://github.com/goodarzilab/iAnalyzer). Briefly, iAnalyzer relies on fitting a generalized linear model to each gene. Coefficients from this GLM were z-score normalized to the negative control guides and finally the largest coefficients were analyzed as potential hits. For the comparison of gene phenotypes between the two cell lines, the DESeq2 ratio of ratios test was used (“DESeq2 Testing Ratio of Ratios (RIP-Seq, CLIP-Seq, Ribosomal Profiling)” n.d.).

### CRISPRi-mediated gene knockdown

Jurkat cells expressing dCas9-KRAB fusion protein were constructed by lentiviral delivery of pMH0006 (Addgene #135448) and FACS isolation of BFP-positive cells.

Guide RNA sequences for CRISPRi-mediated gene knockdown were cloned into pCRISPRia-v2 (Addgene #84832) via BstXI-BlpI sites. After transduction with sgRNA lentivirus, Jurkat cells were selected with 2 µg/mL puromycin (Gibco). The fluorescence of eGFP and of mCherry was measured on BD FACSCelesta Cell Analyzer.

### Reporter cell lines generation

Mutated or wild type sequences of RORC 3’UTR were cloned into the dual GFP-mCherry reporter using MluI-HF and PacI restriction enzymes (NEB) as described above. The reporters were lentivirally dlivered to HEK293 and Jurkat cells and analyzed by flow cytometry as described above.

### Proteasome inhibitor treatment

Jurkat cells were seeded at the density of 0.25*10^7^ cells per mL. Either the proteasome inhibitors (Carfilzonib or Bortezomib, Cayman Chemical) or negative control (DMSO) were added at the given concentration. After 24 hours long incubation, the fluorescence of eGFP and of mCherry was measured on BD FACSCelesta Cell Analyzer.

### T cell isolation, transduction, and Th17 cells differentiation

Th17 cells were derived as described previously (Montoya and Ansel 2017). Plates were coated with 2 µg/mL anti-human CD3 (UCSF monoclonal antibody core, clone: OKT-3) and 4 µg/mL anti-human CD28 (UCSF monoclonal antibody core, clone: 9.3) in PBS with calcium and magnesium for at least 2 h at 37 °C or overnight at 4 °C with plate wrapped in parafilm. Human CD4+ T cells were isolated from human peripheral blood using EasySep human CD4+ T cell isolation kit (17952; STEMCELL) and stimulated in ImmunoCult-XF T cell expansion medium (10981; STEMCELL) supplemented with 10 mM HEPES, 2 mM L-glutamine, 100 µM 2-ME, 1 mM sodium pyruvate, and 10 ng/ml TGF-β. 24 h after T cell isolation and initial stimulation on a 96-well plate, 7 ul of lentivirus was added to each sample. After 24 h, the media was removed from each sample without disturbing the cells and replaced with 200 µl fresh media. After 48 h, cells were stimulated with 1.2 µM ionomycin, 25 nM PMA, and 6 µg/ml brefeldin-A, resuspended by pipetting, incubated for 4 h at 37 °C, and harvested for analysis. Half of each sample was stained for CD4, FoxP3, IL-13, IL-17A, IFN-gamma and analyzed on BD LSRFortessa cell analyzer (see below). The other half of the sample was not stained and was analyzed for the expression of eGFP and mCherry on BD LSRFortessa cell analyzer.

Cultured human T cells were collected, washed, and stained with antibodies against cell surface proteins and transcription factors. Cells were fixed and permeabilized with the eBioscience™ Foxp3 / Transcription Factor Staining Buffer Set or the Transcription Factor Buffer Set (BD Biosciences). Extracellular nonspecific binding was blocked with the anti-CD16/CD32 antibody (clone 2.4G2; UCSF Monoclonal Antibody Core). Intracellular nonspecific binding was blocked with anti-CD16/CD32 Abs and 2% normal rat serum. Dead cells were stained with Fixable Viability Dye eFluor 780 (eBioscience) or Zombie Violet Fixable Viability Kit (BioLegend). Cells were stained with the following fluorochrome-conjugated anti-human Abs: anti-CD4 (Invitrogen 17-0049-42), anti-FOXP3 (eBioscience 25-4777-61), anti-IL-13 (eBioscience 11-7136-41), anti-IL-17A (eBioscience 12-7179-42), and anti-IFNγ (BioLegend 502520). Samples were analyzed on BD LSRFortessa cell analyzer.

### Analysis of capillary electrophoresis data with HiTRACE

Capillary electrophoresis runs from chemical probing and mutate-and-map experiments were analyzed with the HiTRACE MATLAB package (Yoon et al. 2011). Lanes were aligned together, bands fit to Gaussian peaks, background subtracted using the no-modification lane, corrected for signal attenuation, and normalized to the internal hairpin control. The end result of these steps is a numerical array of “reactivity” values for each RNA nucleotide that can be used as weights in structure prediction.

### UPF1 targeted CLIP-seq

Jurkat cells expressing RORC reporters (WT, 77-GA mutant variant or 116-CCCTAAG mutant variant) were harvested and crosslinked by ultraviolet radiation (400 mJ/cm2). Cells were then lysed with low salt wash buffer (1xPBS, 0.1%SDS, 0.5% Sodium Deoxycholate, 0.5% IGEPAL ). To probe preferential UPF1 binding towards different reporters, lysates from 77-GA mutant cells were mixed with lysates from either WT or 116-CCCTAAG mutant cells at a 1:1 ratio prior to immunoprecipitation. Samples were then treated with high (1:3000 RNase A and 1:100 RNase I) and low dose (1:15000 RNase A and 1:500 RNase I) of RNase A and RNase I separately and combined after treatment. To immunoprecipitate UPF1-RNA complex, a UPF1 antibody (Thermo A301-902A) was incubated with Protein A/G beads (Pierce) first and then incubated with the mixed cell lysates for 2 hours at 4 °C. Immunoprecipitated RNA fragments were then dephosphorylated (T4 PNK, NEB), polyadenylated, and end labeled with 3’-Azido-3’-dUTP and IRDye® 800CW DBCO Infrared Dye (LI-COR) on beads. SDS-PAGE was then performed to separate protein-RNA complexes, and RNA fragments were collected from nitrocellulose membrane by proteinase K digestion. cDNA was then synthesized using Takara smarter small RNA sequencing kit reagents with a custom UMI-oligoDT primer (CAAGCAGAAGACGGCATACGAGATNNNNNNNNGTGACTGGAGTTCAGACGTGTGCTCTTCCGATCTTTTTTTTTTTTTTT). RORC reporter locus was then amplified with a custom primer (ACACTCTTTCCCTACACGACGCTCTTCCGATCT TGGGGTGATCCAAATACCACC) and sequencing libraries were then prepared with SeqAmp DNA Polymerase (Takara). Libraries were then sequenced on an illumina Hiseq 4000 sequencer.

## Acknowledgements

The authors thank Christopher Mathy, Andrew Natale, Maxim Imakaev, Yessica Gomez for helpful discussions. HG is an Era of Hope Scholar (W81XWH-2210121) and supported by R01CA240984 and R01CA244634. This work was partly supported by National Institutes of Health (NIH) grants 1R35GM140847 (Y.C.). L.A.G. is funded by an NIH New Innovator Award (DP2 CA239597), a Pew-Stewart Scholars for Cancer Research award and the Goldberg-Benioff Endowed Professorship in Prostate Cancer Translational Biology. Cryo-EM equipment at UCSF is partially supported by NIH grants S10OD020054, S10OD021741 and S10OD026881. YC is an Investigator of Howard Hughes Medical Institute. Sequencing was performed at the UCSF CAT, supported by UCSF PBBR, RRP IMIA and NIH 1S10OD028511-01 grants. A.N. was supported by DoD PRCRP Horizon Award W81XWH-19-1-0594. L.F. was supported by NIH training grant T32CA108462-15.

## Contributions

M.K. and H.G. designed the study. M.K. developed SwitchFinder. M.K. and A.N. performed massively parallel reporter assays. M.K. and L.F. performed DMS-MaPseq experiments. M.K. and C.P. performed SHAPE experiments. D.A. and Y.C. performed cryogenic electron microscopy experiments. M.K. performed mutagenesis experiments. M.K. performed antisense oligo transfection experiments. M.K., S.K.Z. and K.M.A performed Th17 differentiation experiments. M.K, A.W. and L.G. performed CRISPRi screens. M.K. performed CRISPRi knockdown experiments. M.K. and J.Y. performed proteasome inhibition experiments. S.Z. performed CLIP-seq experiments. M.K. and H.G. wrote the manuscript with input from all authors.

## Competing interests

M.K. and H.G. are inventors on a provisional patent related to this study. L.A.G has filed patents on CRISPR functional genomics.

## Ethics statement

This research complies with all relevant ethical regulations

## Data and code availability

Sequencing data has been deposited in the Gene Expression Omnibus (GEO). SwitchFinder source code is available at https://github.com/goodarzilab/SwitchFinder

## Notes

### Competing Interest Statement

The authors have declared no competing interest.

