## Supplementary Figures for "A systematic search for RNA structural switches across the human transcriptome"

#### The PDF file includes:

Suppl. Fig. 1: SwFinder identifies saddles in RNA folding energy landscape
Suppl. Fig. 2: Overview of high-throughput screening approaches for improved RNA switch predictions
Suppl. Fig. 3: In vitro SHAPE reactivity of the RORC RNA switch sequence in vitro.
Suppl. Fig. 4: CryoEM reveals distinct clusters of RNA structures
Suppl. Fig. 5: Differentiation of Th17 cells from primary human CD4+ cells
Suppl. Fig. 6: CRISPRi screen highlights the pathways acting downstream of the RORC RNA switch

#### Other Supplementary Material for this manuscript includes the following:

Data file S1 (Microsoft Excel format). AUC values for SwFinder prediction of RNA switches from common riboswitch Rfam families
Data file S2 (Microsoft Excel format). SwFinder predictions for the 3,750 RNA fragments selected for further in vivo screening
Data file S3 (Microsoft Excel format). mRNA and gDNA read counts in the sorted bins of the functional screen
Data file S4 (Microsoft Excel format). DMS-MaPseq accessibility profiles and the second iteration of SwFinder predictions for 1,454 high confidence RNA switches
Data file S5 (Microsoft Excel format). mRNA and gDNA read counts in the sorted bins for the massively parallel mutagenesis analysis
Data file S6 (Microsoft Excel format). Sequences of RORC mutant sequences referred to in the paper
Data file S7 (Microsoft Excel format). Cryo-EM data collection, refinement and validation statistics
Data file S8 (Microsoft Excel format). Cryo-EM data collection parameters
Data file S9 (Microsoft Excel format). Antisense oligonucleotides (ASO) used for RORC structural ensemble perturbation
Data file S10 (Microsoft Excel format). gDNA read counts for the CRISPRi screens

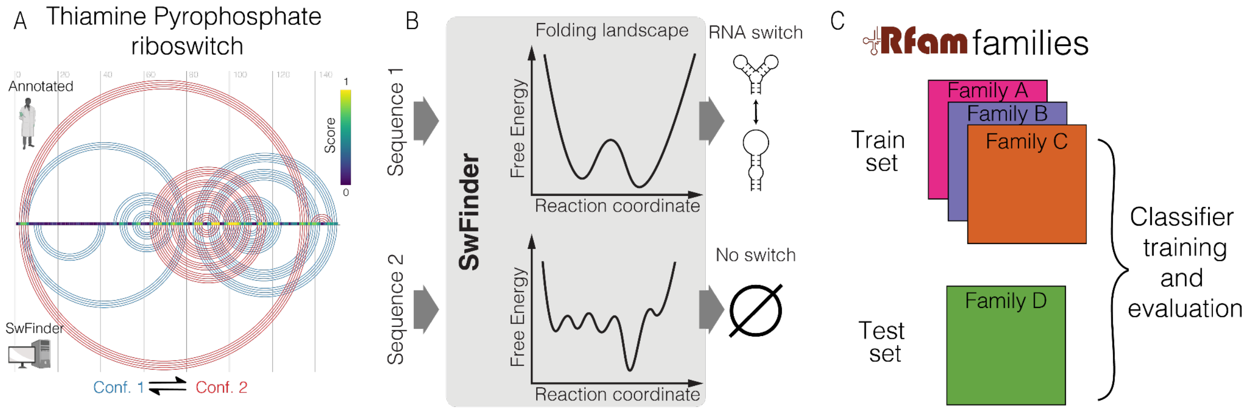


**Suppl. Fig. 1: SwFinder identifies saddles in RNA folding energy landscape**

(A) Example of SwFinder locating the thiamine pyrophosphate RNA switches within the mRNA sequence. Top: arc representation of the RNA base pairs that change between the two conformations of the RNA switch, as in (Barsacchi et al. 2016). The two conformations are shown in red and blue, respectively. Bottom: the two conformations of the RNA switch as predicted by SwFinder. Middle: SwFinder score reflecting the likelihood of a given nucleotide to be involved in two mutually exclusive base pairings.

(B) Scheme of SwFinder model. SwFinder analyzes RNA folding energy landscape of a given RNA sequence and assigns higher score to the landscapes that demonstrate riboswitch-like features.

(C) The setup for evaluating the ability of a model to find RNA switches from novel families. At the classifier training step, riboswitches from one of the RFAM families get separated into the “test set”, while the model gets trained on the riboswitches from other RFAM families. The test set then is used to evaluate the model performance

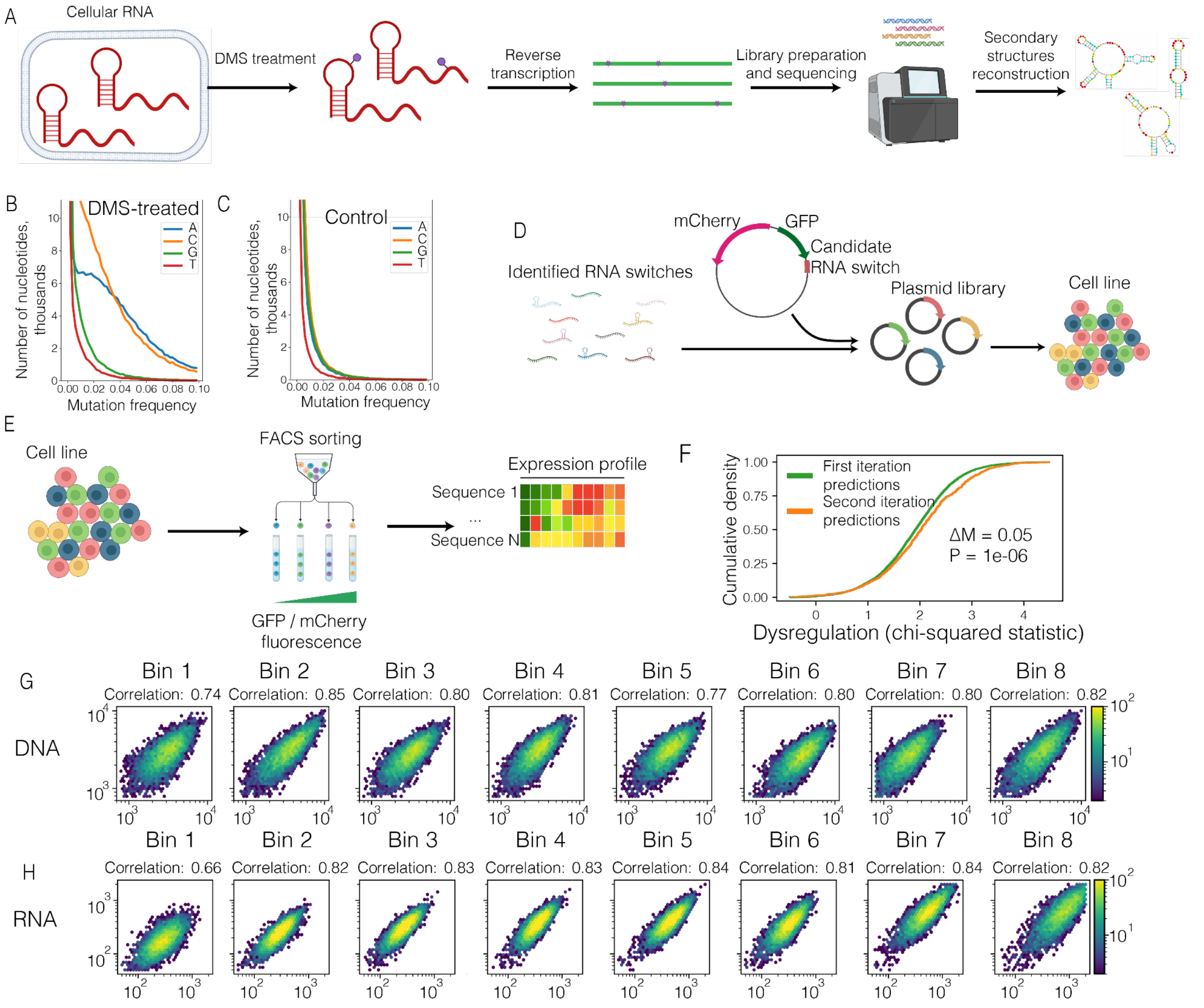


**Suppl. Fig. 2: Overview of high-throughput screening approaches for improved RNA switch predictions**

(A) Overview of DMS-MaPseq workflow. Mammalian cells are treated with DMS. DMS-modified nucleotides cause mutations when cDNA is synthesized from RNA templates. The cDNA libraries are sequenced, the DMS-caused mutations are counted, providing the flexibility estimates for each A- or C- nucleotide.

(B) Cumulative mutation frequency in DMS-treated candidate riboswitches, separated by nucleotide.

(C) Cumulative mutation frequency in non-treated candidate riboswitches, separated by nucleotide.

(D) Overview of the library generation workflow for Massively Parallel Reporter Assay (MPRA). Sequences of candidate RNA switches are synthesized as DNA oligonucleotides and cloned into a reporter vector into 3'UTR region of a eGFP cDNA. The plasmid library is packaged into lentiviral particles, and used for infecting mammalian cells. The infection is performed at low MOI (infection rate) to ensure that most cells get only a single plasmid copy.

(E) Overview of the MPRA workflow. A population of mammalian cells is separated into bins based on GFP/mCherry fluorescence ratio. In the schematic, cells are colored according to the sequence they carry in the 3'UTR of the GFP reporter

(F) Cumulative density plot of dysregulation values, comparing the candidate RNA switches predicted in first and second (DMS-MaPseq informed) iterations of SwFinder. Dysregulation values are estimated using chi-square test for every individual candidate RNA switch across 8 expression bins. Median difference ($\Delta$M) and p value (calculated using Mann-Whitney U-test) are shown.

(G) Correlations of read counts of gDNA libraries between the biological replicates of massively parallel mutagenesis analysis

(H) Correlations of read counts of RNA libraries between the biological replicates of massively parallel mutagenesis analysis

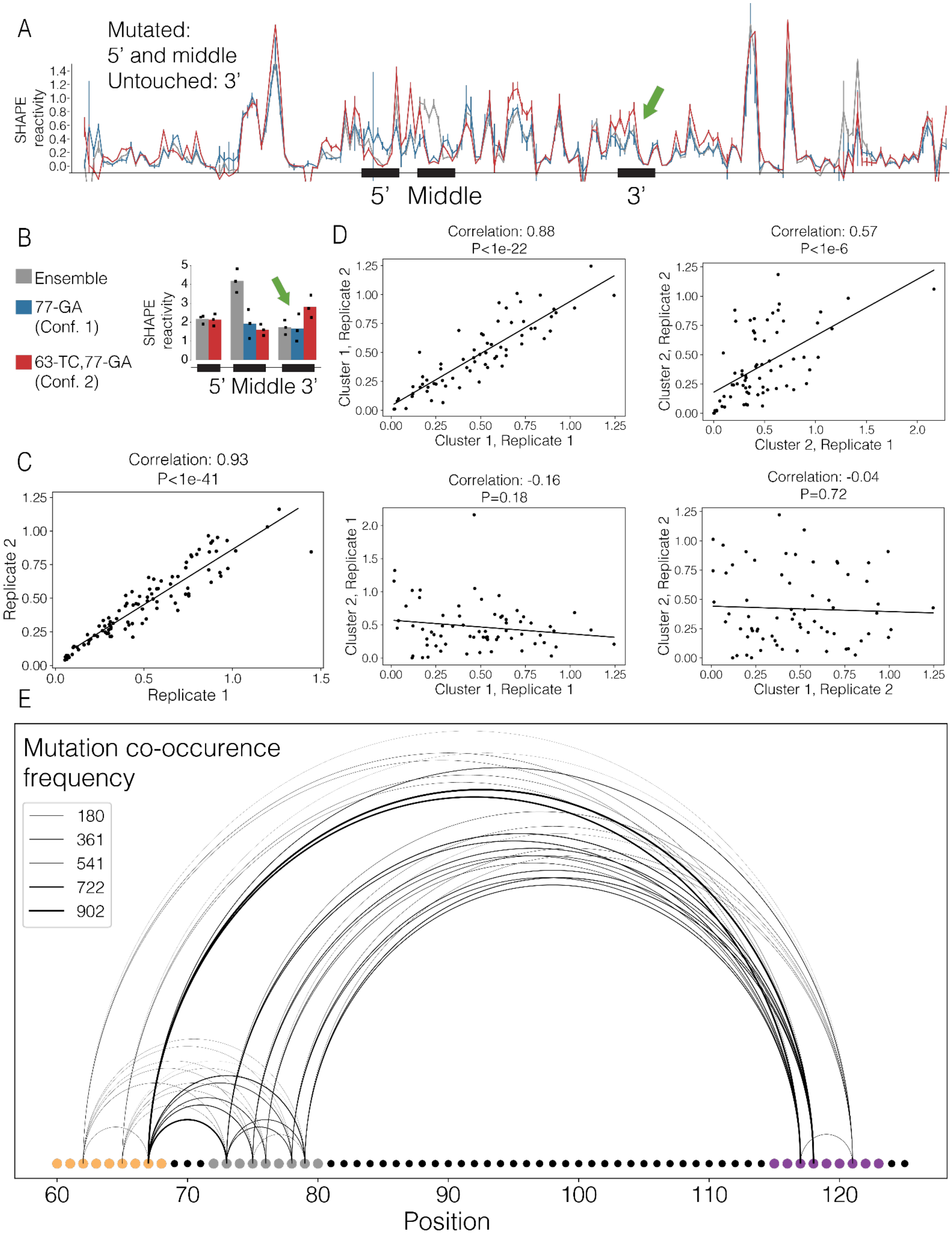


**Suppl. Fig. 3: In vitro SHAPE reactivity of the RORC RNA switch sequence *in vitro*.**

(A) SHAPE reactivity profiles for the wild type sequence and for the mutation-rescue pair of sequences (blue - "77-GA", red - "63-TC,77-GA"). The three switching regions are labeled. The SHAPE reactivity changes in the non-mutated regions are highlighted in bold arrows. N replicates = 3.

(B) Barplots of cumulative SHAPE reactivity within the switching regions for the wild type sequence (in gray) and for the mutation-rescue pair of sequences (blue - “77-GA”, red - “63-TC,77-GA). N replicates = 3.

(C) Scatter plot showing the reproducibility of the DMS signal between two replicates. Each dot represents a single nucleotide. Normalized DMS signal is shown on both axes.

(D) Scatter plots showing the reproducibility of the DMS signal within each conformation between two replicates. Each dot represents a single nucleotide. Top left: normalized DMS signal of the conformation 1 is shown on both axes, for different replicates. Top right: normalized DMS signal of the conformation 2 is shown on both axes, for different replicates. Bottom left: normalized DMS signal of the conformations 1 and 2 is shown on the two axes, respectively, within the replicate 1. Bottom right: normalized DMS signal of the conformations 1 and 2 is shown on the two axes, respectively, within the replicate 2.

(E) Arc plot showing the co-occurence of mutations in given pairs of nucleotides on the same read. The nucleotides of the switching regions are shown, and the switching regions are color-coded: the 5' region is shown in orange, the middle region - in gray, and the 3’ region - in purple. The thickness of the arc is proportional to the number of reads where the mutations of two nucleotides co-occur.

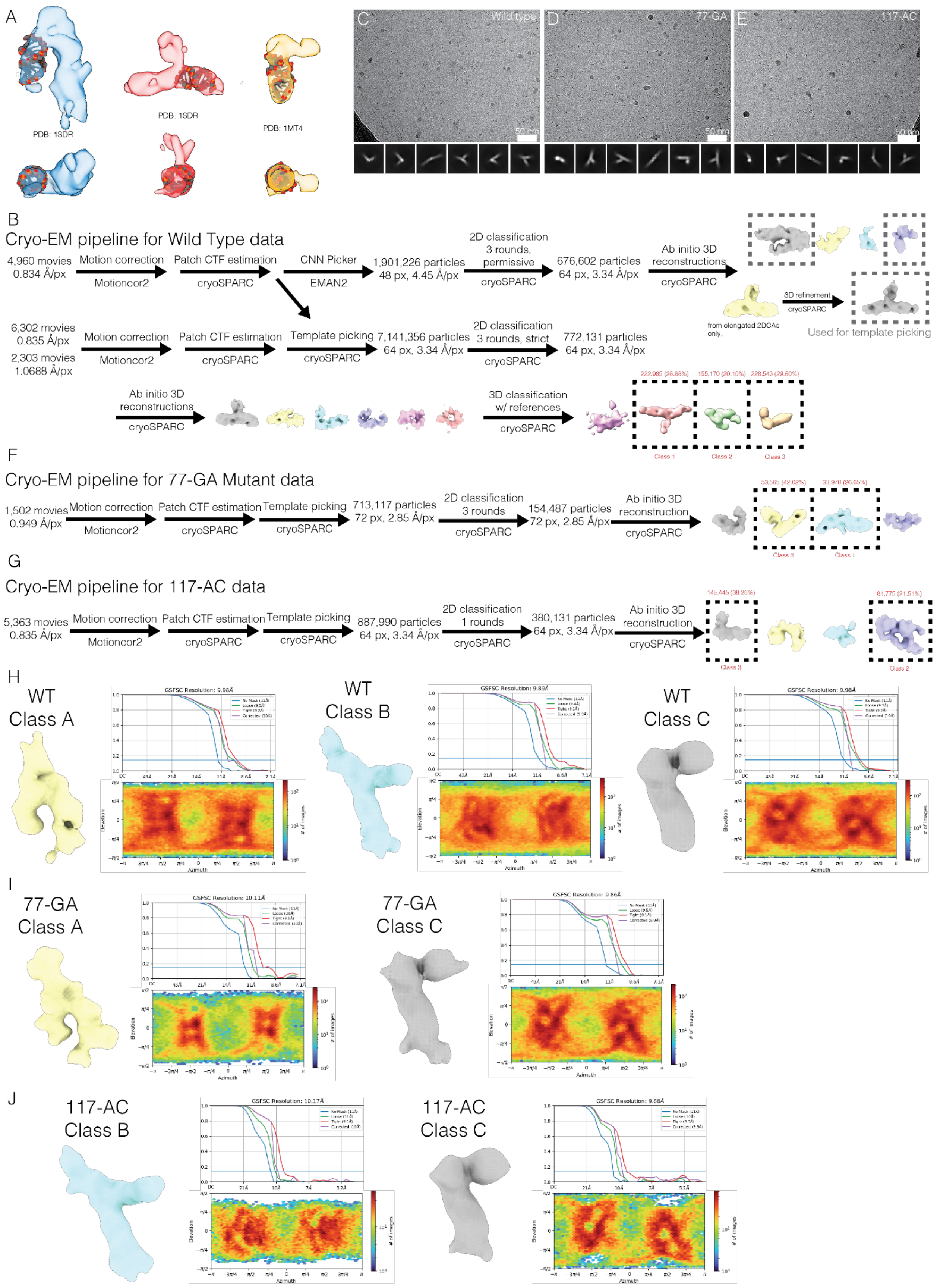
A
PDB: 1SDR
PDB: 1SDR
PDB: 1MT4
C
Wild type
D
77-GA
E
117-AC
50 nm
B
Cryo-EM pipeline for Wild Type data
4,960 movies
0.834 Å/px
Motion correction
Motioncor2
Patch CTF estimation
cryoSPARC
CNN Picker
EMAN2
1,901,226 particles
48 px, 4.45 Å/px
2D classification
3 rounds,
permissive
cryoSPARC
676,602 particles
64 px, 3.34 Å/px
Ab initio 3D
reconstructions
cryoSPARC
3D refinement
cryoSPARC
from elongated 2DCAs only.
Used for template picking
6,302 movies
0.835 Å/px
2,303 movies
1.0688 Å/px
Motion correction
Motioncor2
Patch CTF estimation
cryoSPARC
Template picking
cryoSPARC
7,141,356 particles
64 px, 3.34 Å/px
2D classification
3 rounds, strict
cryoSPARC
772,131 particles
64 px, 3.34 Å/px
Ab initio 3D
reconstructions
cryoSPARC
3D classification
w/ references
cryoSPARC
222,985 (28.88%)
155,170 (20.10%)
228,543 (29.60%)
Class 1
Class 2
Class 3
F
Cryo-EM pipeline for 77-GA Mutant data
1,502 movies
0.949 Å/px
Motion correction
Motioncor2
Patch CTF estimation
cryoSPARC
Template picking
cryoSPARC
713,117 particles
72 px, 2.85 Å/px
2D classification
3 rounds
cryoSPARC
154,487 particles
72 px, 2.85 Å/px
Ab initio 3D
reconstruction
cryoSPARC
Class 3
Class 1
G
Cryo-EM pipeline for 117-AC data
5,363 movies
0.835 Å/px
Motion correction
Motioncor2
Patch CTF estimation
cryoSPARC
Template picking
cryoSPARC
887,990 particles
64 px, 3.34 Å/px
2D classification
1 rounds
cryoSPARC
380,131 particles
64 px, 3.34 Å/px
Ab initio 3D
reconstruction
cryoSPARC
Class 3
Class 2
H
WT
Class A
GSFSC Resolution: 9.98Å
WT
Class B
GSFSC Resolution: 9.89Å
WT
Class C
GSFSC Resolution: 9.98Å
Elevation
Azimuth
### of images
I
77-GA
Class A
GSFSC Resolution: 10.11Å
77-GA
Class C
GSFSC Resolution: 9.86Å
J
117-AC
Class B
GSFSC Resolution: 10.17Å
117-AC
Class C
GSFSC Resolution: 9.88Å

**Suppl. Fig. 4: CryoEM reveals distinct clusters of RNA structures**
(A) Class A (left), B (middle), and C (right) of WT RORC RNA overlaid with stereotypical RNA tertiary structures from PDB including dsRNA B-helix and RNA hairpin. Features representing the major groove and a hairpin are visible in regions of the maps.
(B) Schematic cryo-EM image processing pipelines for WT RORC RNA. During template picking, templates and micrographs were low-pass filtered to 20 Å.
(C-E) Representative micrographs and 2D class averages for RORC RNA switch WT sequence (C), 77-GA (D) and 117-AC (E).
(F,G) Schematic cryo-EM image processing pipelines for 77-GA (F), and 117-AC (G) mutants. During template picking, templates and micrographs were low-pass filtered to 20 Å.
(H) Gold-standard half-map refinement volume, FSC curves, and orientation distribution plot for 3D classes from WT RNA sample.
(I) Gold-standard half-map refinement volume, FSC curves, and orientation distribution plot for 3D classes from 77-GA sample.
(J) Gold-standard half-map refinement volume, FSC curves, and orientation distribution plot for 3D classes from 117-AC sample.

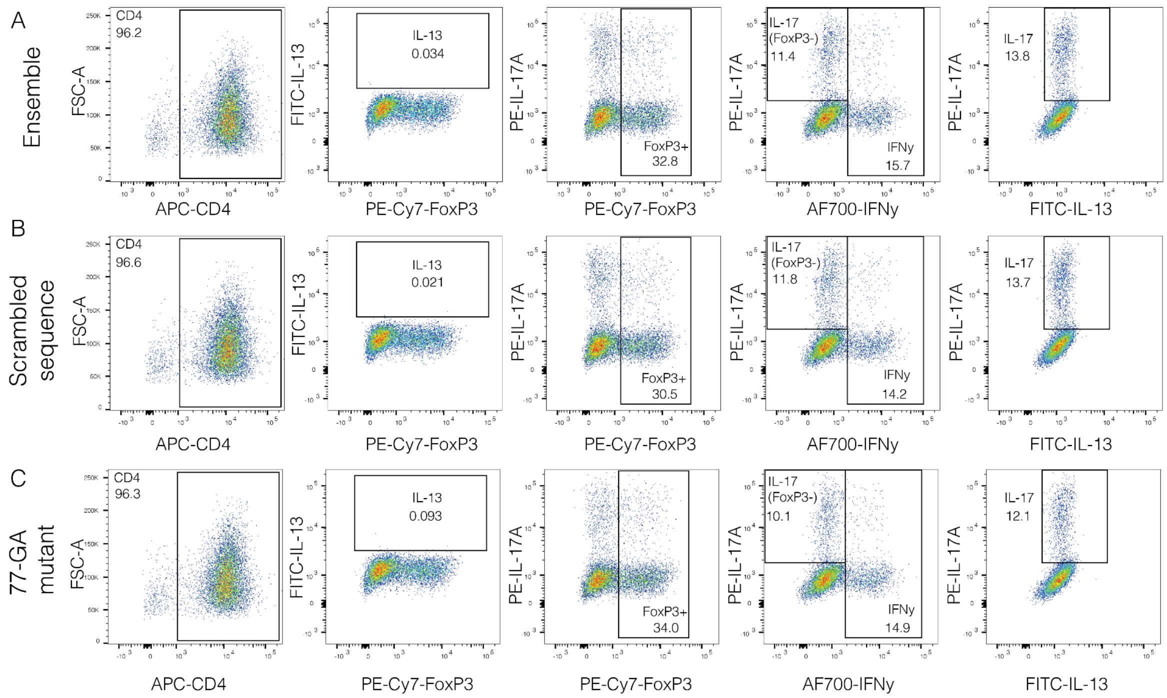


**Suppl. Fig. 5: Differentiation of Th17 cells from primary human CD4+ cells**
Representative fluorescence-activated cell sorting plots of CD4+ T cells for the same samples as in Fig. 5D. On the day 5 of differentiation, each sample was split in half; one half was analyzed for mCherry and GFP expression (shown in Fig. 5D), the other half was stained for the expression of CD4, FoxP3, IL-13, IL-117A, IFN-gamma. The cells expressing a given marker are highlighted with a frame and a fraction of the parental cellular population is given. Each sample was analyzed in 4 replicates; a single representative replicate is displayed for each sample. The cells express the reporter construct containing: (A) wild type RORC 3'UTR sequence, (B) scrambled RORC RNA switch, (C) "77-GA" mutant

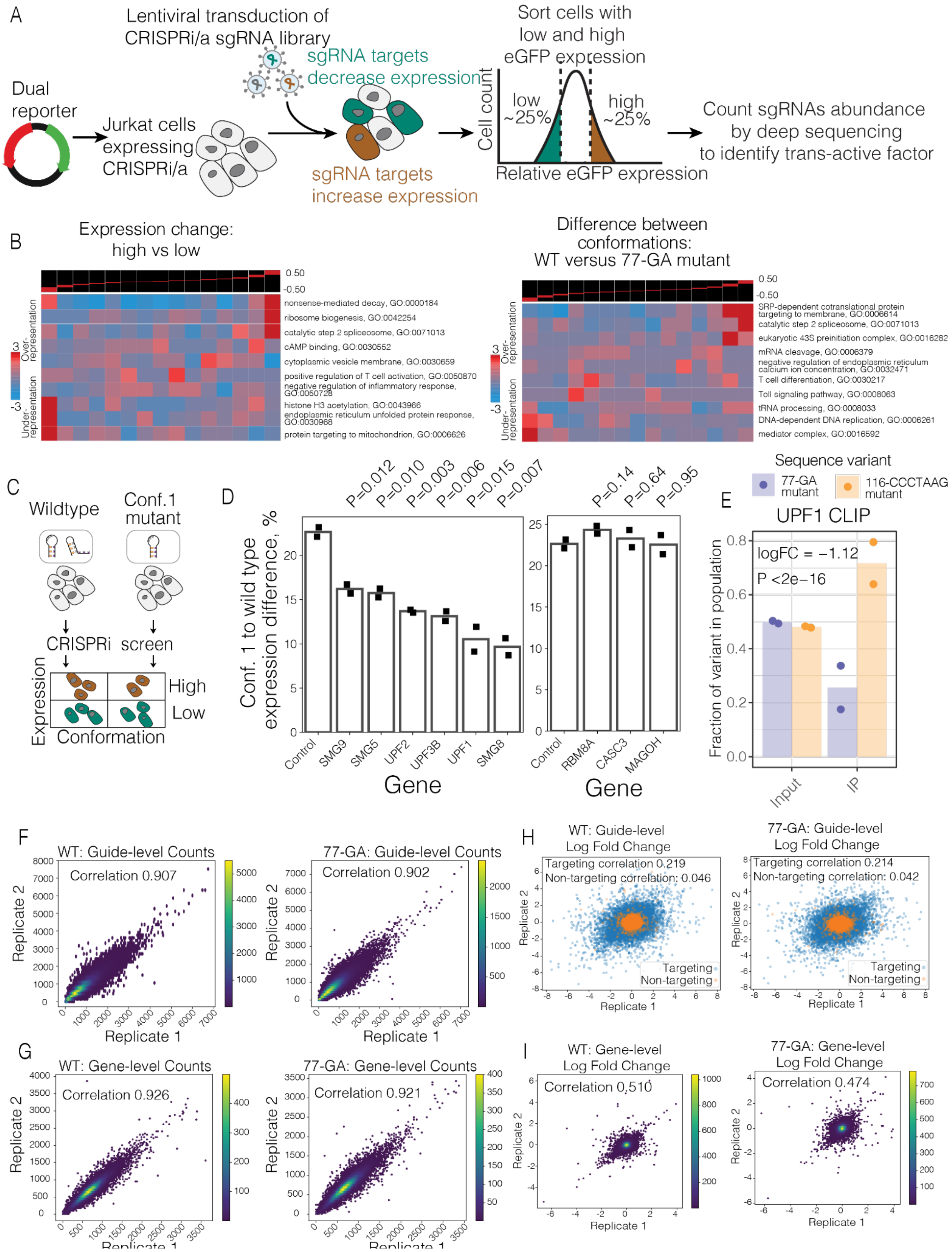


**Suppl. Fig. 6: CRISPRi screen highlights the pathways acting downstream of the RORC RNA switch**

(A) Overview of the flow cytometry-based CRISPRi screen workflow. Dual reporter construct expressing RORC RNA switch sequence in the 3'UTR of eGFP gene is stably expressed by CRISPRi-active Jurkat cells. A genome-wide library of sgRNAs (Horlbeck et al., 2018) is then delivered to the cell line expressing the reporter construct. Cells are separated into individual bins based on relative fluorescence levels of eGFP and mCherry. gDNA is extracted from individual bins and sequenced; the relative abundance of sgRNAs in individual bins is counted.

(B) Gene-set enrichment analysis of the data depicted in Fig. 6A (left) and Fig. 6B (right). The genes were distributed into equally populated bins based on their comparative abundance between high expression and low expression quartiles (left), or based on their comparative phenotype in the CRISPRi screens performed in WT or 77-GA mutant backgrounds (right). Then the enrichment of a given gene set was calculated in each bin using iPAGE, a mutual information-based algorithm (Goodarzi et al. 2009).

(C) Experiment design table. Columns: two Jurkat cell lines were analyzed, each stably expressing a reporter construct with RORC RNA switch sequence located in the reporter gene 3'UTR. One cell line ("WT") expressed the wild type sequence of the RORC RNA switch, the other cell line ("77-GA mutant") expressed a mutated sequence that locks the RNA switch in the conformation 1. Rows: for each cell line, the top and the bottom quantiles of the expression distribution were taken (see the panel (A).

(D) The effect of knockdown of SURF and EJC complex member proteins on the expression change upon the conformation equilibrium shift. The individual genes were knocked down using the CRISPRi system in both WT and 77-GA mutant cell lines, then the change of reporter gene expression was measured by flow cytometry (N replicates = 2). The bar plots demonstrate the expression ratios of WT to 77-GA mutation cell lines.

(E) The fractions of reads carrying the 77-GA mutant sequence or 116-CCCTAAG mutant sequence in UPF1 cross-linking and immunoprecipitation (CLIP) library. Left: input RNA libraries, extracted from the 116-CCCTAAG and 77-GA mutant expressing Jurkat cells, mixed at 1:1 ratio. Right: libraries after anti-UPF1 immunoprecipitation

(F) Density plots showing the correlation of sgRNA counts between the replicates of the CRISPRi screens performed in the WT (left) and 77-GA mutant (right) backgrounds.

(G) Density plots showing the correlation of gene counts between the replicates of the CRISPRi screens performed in the WT (left) and 77-GA mutant (right) backgrounds. The counts of all the sgRNAs targeting a given gene are pooled and reported as a single number (N = 5 sgRNAs per gene).

(H) Scatter plots showing the correlation of sgRNA phenotypes between the replicates of the CRISPRi screens performed in the WT (left) and 77-GA mutant (right) backgrounds. Logarithmic fold changes between the sgRNA abundance "high" and "low" expression bins (measured relative fluorescence levels of eGFP and mCherry) are shown on both axes. Non-targeting sgRNAs are shown in orange; all the other sgRNAs are shown in blue. The correlation values are reported separately for non-targeting and targeting sgRNAs.

(I) Density plots showing the correlation of gene phenotypes between the replicates of the CRISPRi screens performed in the WT (left) and 77-GA mutant (right) backgrounds. Logarithmic fold changes between the abundance of sgRNAs targeting a given gene in “high” and “low” expression bins (measured relative fluorescence levels of eGFP and mCherry) are shown on both axes.
